# Hemato-vascular specification requires *arnt1* and *arnt2* genes in zebrafish embryos

**DOI:** 10.1101/2022.01.04.474920

**Authors:** Hailey E. Edwards, Mary Jane Edgington, Jaclyn P. Souder, Daniel A. Gorelick

## Abstract

During embryonic development, a subset of cells in the mesoderm germ layer are specified as hemato- vascular progenitor cells, which then differentiate into endothelial cells and hematopoietic stem and progenitor cells. In zebrafish, the transcription factor *npas4l*, also known as *cloche*, is required for the specification of hemato-vascular progenitor cells. However, it is unclear if *npas4l* is the sole factor at the top of the hemato-vascular specification cascade. Here we show that *arnt1* and *arnt2* genes are required for hemato-vascular specification. We found that *arnt1;arnt2* double homozygous mutant zebrafish embryos (herein called *arnt1/2* mutants), but not *arnt1* or *arnt2* single mutants, lack blood cells and most vascular endothelial cells. *arnt1/2* mutants have reduced or absent expression of *etsrp* and *tal1*, the earliest known endothelial and hematopoietic transcription factor genes. *npas4l* and the *arnt* genes are PAS domain-containing bHLH transcription factors that function as dimers. We found that Npas4l binds both Arnt1 and Arnt2 proteins *in vitro*, consistent with the idea that PAS domain- containing bHLH transcription factors act in a multimeric complex to regulate gene expression. Our results demonstrate that *npas4l*, *arnt1* and *arnt2* act together to regulate endothelial and hematopoietic cell fate, where each gene is necessary, but by itself not sufficient, to drive hemato-vascular specification. Our results also demonstrate that *arnt1* and *arnt2* act redundantly in a transcriptional complex containing *npas4l*, but do not act redundantly when interacting with another PAS domain- containing bHLH transcription factor, the aryl hydrocarbon receptor. Altogether, our data enhance our understanding of hemato-vascular specification and the function of PAS domain-containing bHLH transcription factors.

## INTRODUCTION

Endothelial and hematopoietic cells are derived from a common mesodermal progenitor (Huber et al., 2004; Kataoka et al., 2011; Liao et al., 2000). In mice and zebrafish, fully multipotent hematopoietic stem cells (HSCs) are derived from specialized endothelial cells, called hemogenic endothelium (Hirschi, 2012; Kobayashi et al., 2014; Ramírez-Bergeron et al., 2004) The first hemato-vascular progenitors were traced back to the lateral plate mesoderm of zebrafish and in the extraembryonic yolk sac mesoderm of mice (Lee et al., 1994; Palis et al., 1999; Stainier et al., 1993). The complete mechanism of the specification of these hemato-vascular progenitors from the multipotent mesoderm remains elusive in mammals.

Many genes involved in embryonic hematopoiesis and vascular development are evolutionally conserved between zebrafish and mammals like mice and humans (Begley et al., 1989; Casie Chetty et al., 2017; Dooley et al., 2005; Ferrara et al., 1996; Herbert and Stainier, 2011; Kalev-Zylinska et al., 2002; Kataoka et al., 2011; Okuda et al., 1996; Wang et al., 1996). In zebrafish, the *neuronal PAS4-like* gene (*npas4l*), formerly known as *cloche,* is the earliest known transcription factor to drive the specification of hemato-vascular progenitors (Reischauer et al., 2016). *npas4l* mutants have a distinctive phenotype as they lack most endothelial and all hematopoietic cells (Stainier et al., 1995; Thompson et al., 1998).

Npas4l is a basic helix-loop-helix Per-Arnt-Sim (bHLH-PAS) transcription factor and is thought to be the master regulator of hemato-vascular specification (Reischauer et al., 2016). bHLH-PAS transcription factors are classified as Class I or Class II, which must dimerize to regulate gene transcription or repression (**Figure 1A**) (Card et al., 2005; Edwards and Gorelick, 2022; Pongratz et al., 1998). Npas4l is a Class I bHLH-PAS protein, but its Class II dimerization partner during hemato-vascular progenitor specification is not known. We hypothesized that Arnt1 or Arnt2 is the Class II dimerization partner of Npas4l in zebrafish because *arnt1* and *arnt2* genes are expressed at the same stages of embryo development as hemato-vascular specification (Baranasic et al., 2022; Prasch et al., 2006; Reischauer et al., 2016; Tanguay et al., 2000).

**Figure 1.**
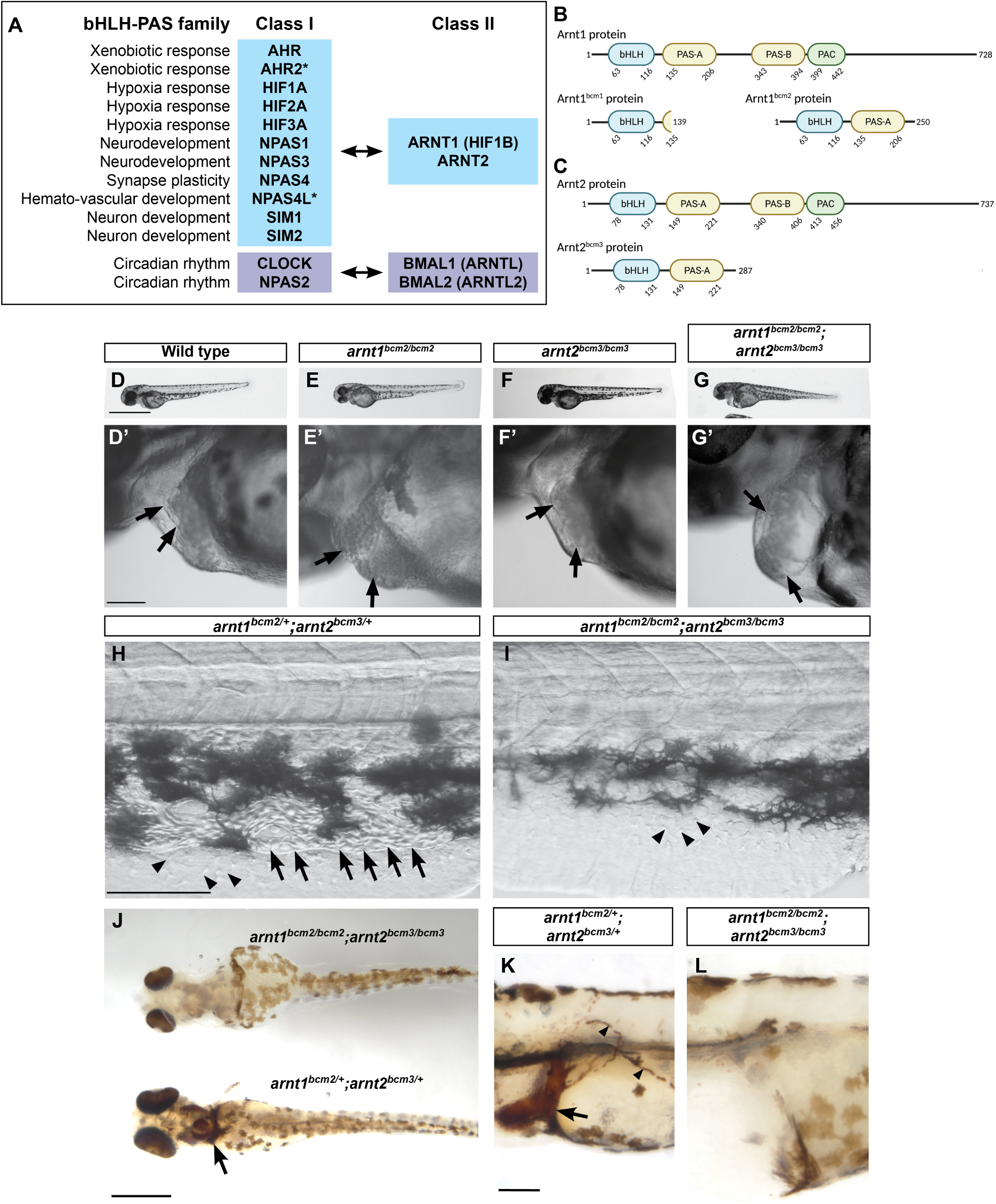
*arnt1*/2 double mutant embryos, but not *arnt1* or *arnt2 single* mutants, lack circulating blood, have cardiac edema and atrial enlargement. (A) bHLH-PAS transcription factors act as heterodimers: a Class I protein and Class II protein interact and bind DNA. Asterisks indicate proteins encoded by genes present in zebrafish but absent in humans. (B,C) Generation of zebrafish with mutations in *arnt1* and *arnt2* genes. The amino acid sequences for the zebrafish Arnt1 (B), and Arnt2 proteins (C) are represented by the black line. Basic helix-loop-helix (bHLH), Per-Arnt-Sim (PAS) domains, and the motif C-terminally located to the PAS (PAC) are represented by the labeled bubbles. The numbers below each domain indicate the starting and ending amino acid (AA) for each motif. (B) The *arnt1^bcm1^*mutation causes a premature stop codon to occur at AA139 and loss of the PAS-B, PAC, and majority of the PAS-A. The *arnt1^bcm2^* mutation causes a premature stop codon to occur at AA250 and loss of the PAS-B and PAC domains. (C) The *arnt2^bcm3^* mutation causes a premature stop codon to occur at AA287 and loss of the PAS-B and PAC domains. (D-I) Live images of embryos from the indicated genotype at 2 days post fertilization (dpf). All double homozygous *arnt1^bcm2/bcm2^; arnt2^bcm3/bcm3^* embryos showed cardiac edema and atrial enlargement (G and G’). The wildtype (D and D’), *arnt1^bcm2/bcm2^* (E and E’), and *arnt2^bcm3/bcm3^* (F and F’) siblings of the double homozygous mutants (*arnt1^bcm2/bcm2^; arnt2^bcm3/bcm3^*) showed normal heart development. Black arrows in panels D’-G’ indicate the atrium and ventricle of the heart. Images are representative of 7 different clutches, each clutch contained 20-300 embryos. All siblings of double homozygous mutants (*arnt1^bcm2/bcm2^; arnt2^bcm3/bcm3^*) had circulating blood (H and Movie 1, Movie 2, Movie 5, Movie 6) while *arnt1/2* mutants lacked visible circulating blood cells (I and Movie 3, Movie 4, Movie 7, Movie 8). The black arrows in panel H indicate blood cells. The arrowheads in H and I indicate non-blood cells found in all embryos. (J-L), o-dianisidine staining to label blood cells in embyos 3 days post fertilization shows that *arnt1^bcm2/bcm2^*; *arnt2^bcm3/bcm3^* double homozygous mutants lack blood cells, while all other genotypes have clearly labeled blood cells. The arrow in J and K points to the duct of Cuvier, and the arrowheads in K show circulating blood cells. All embryos were derived from *arnt1^bcm2/+^;arnt2^bcm3/+^* parents, images are representative of 1 clutch, containing 98 embryos. (J) N=96 of 98 embryos showed clear o-dianisidine staining and none of these embryos were *arnt1/2* mutants. (K) N=2 of 98 embryos showed a lack of o-dianisidine staining and both embryos were confirmed *arnt1/2* mutants. For all images anterior is towards the left and dorsal is towards the top, except for panel J which shows a ventral view of the embryos. Scale bars: D-G, 1 mm; D’-G’ and H-K, 100 μm.

To test this hypothesis, we examined hemato-vascular development in zebrafish with predicted loss-of-function mutations in *arnt1* and *arnt2* genes. We find that Arnt1 and Arnt2 are essential for hemato-vascular specification. Double homozygous *arnt1;arnt2* zebrafish embryos (herein called *arnt1/2* mutants) phenocopy *npas4l* mutants and show lack of circulating blood, absence of hematopoietic stem and progenitor cells, and dramatic reduction in the number of vascular endothelial cells. We find that expression of *npas4l*-target genes requires *npas4l* and *arnt1* or *arnt2*. Additionally, we show that zebrafish Npas4l protein interacts with both zebrafish Arnt1 and zebrafish Arnt2 proteins *in vitro*. Together, our results suggest that Arnt1 and Arnt2 act redundantly and form a transcriptional complex with Npas4l that sits atop the hemato-vascular specification cascade.

## MATERIALS AND METHODS

### Zebrafish

Adult zebrafish were raised at 28.5°C on a 14-h light, 10-h dark cycle in the BCM Zebrafish Research Facility in an Aquaneering recirculating water system (Aquaneering, Inc., San Diego, CA) and a Tecniplast recirculating water system (Tecniplast USA, Inc., West Chester, PA). Wild-type zebrafish were AB strain (Westerfield, 2000) and *arnt1* and *arnt2* mutant lines were generated on the AB strain. Once the mutant lines were established, they were bred onto the *Tg(Fli1:EGFP)^y1^* transgenic background (Lawson and Weinstein, 2002). All procedures were performed in accordance with and approved by the BCM Institutional Animal Care and Use Committee.

### Embryo collection

Adult zebrafish were allowed to spawn naturally in pairs or in groups. Embryos were collected in intervals of 20 minutes to ensure precise developmental timing or staged following collection, placed in 60 cm^2^ Petri dishes at a density of no more than 100 per dish in E3B media (60X E3B: 17.2g NaCl, 0.76g KCl, 2.9g CaCl2-2H2O, 2.39g MgSO4 dissolved in 1L Milli-Q water; diluted to 1X in 9L Milli-Q water plus 100 μL 0.02% methylene blue), and then stored in an incubator at 28.5°C on a 14-h light, 10-h dark cycle.

### Genotyping adult zebrafish

Genomic DNA was isolated from tail biopsies from individual adult zebrafish and incubated in 50 μL ELB (10 mM Tris pH 8.3, 50 mM KCl, 0.3% Tween 20) with 0.5 μL proteinase K (800 U/ml, NEB) in 96 well plates, one sample per well, at 55°C for 8 hours. Proteinase K was inactivated by incubation at 98°C for 10 minutes and DNA was stored at -20°C. Genotyping was performed by PCR and high- resolution melting curve analysis (HRMA) as described (Parant et al., 2009; Romano et al., 2017). All melting curves were generated with a Bio-Rad CFX96 or CFX Opus 96 Real-Time System over a 70- 95°C range and analyzed with the Bio-Rad CFX Manager 3.1 or the Bio-Rad CFX Maestro 4.1 software. All mutations were confirmed by TA cloning and sequencing. All primers used for HRMA can be found in **Table S1**.

**Table 1.** Genotypes of embryos from arnt1^bcm2/+^; arnt2^bcm3/+^ parents follow expected Mendelian ratios. The frequencies of the observed genotypes were not significantly different from the expected genotypes, X^2^ (8, N=356) = 4.38, p>0.05.

| **Genotype** | **Expected**<br>**Percentage** | **Expected**<br>**n=356** | **Actual**<br>**Percentage** | **Actual**<br>**n=356** |
| --- | --- | --- | --- | --- |
| *arnt1*<sup>+/+</sup>; *arnt2*<sup>+/+</sup> | 6.3% | 22.25 | 6.2% | 22 |
| *arnt1*<sup>+/+</sup>; *arnt2*<sup>bcm3/+</sup> | 12.5% | 44.5 | 14.3% | 51 |
| *arnt1*<sup>+/+</sup>; *arnt2*<sup>bcm3/bcm3</sup> | 6.3% | 22.25 | 4.5% | 16 |
| *arnt1*<sup>bcm2/+</sup>; *arnt2*<sup>+/+</sup> | 12.5% | 44.5 | 11.8% | 42 |
| *arnt1*<sup>bcm2/+</sup>; *arnt2*<sup>bcm3/+</sup> | 25% | 89 | 25.3% | 90 |
| *arnt1*<sup>bcm2/+</sup>; *arnt2*<sup>bcm3/bcm3</sup> | 12.5% | 44.5 | 11.8% | 42 |
| *arnt1*<sup>bcm2/bcm2</sup>; *arnt2*<sup>+/+</sup> | 6.3% | 22.25 | 5.3% | 19 |
| *arnt1*<sup>bcm2/bcm2</sup>; *arnt2*<sup>bcm3/+</sup> | 12.5% | 44.5 | 13.5% | 48 |
| *arnt1*<sup>bcm2/bcm2</sup>; *arnt2*<sup>bcm3/bcm3</sup> | 6.3% | 22.25 | 7.3% | 26 |

### CRISPR-Cas9 Mutant Generation

Cas9 mRNA and gRNAs for *arnt1* and *arnt2* mutants were generated as previously described (Romano et al., 2017; Souder and Gorelick, 2019). Cas9 mRNA was transcribed from a linearized pT3TS-nCas9n plasmid, Addgene #46757 (Jao et al., 2013). The target sequences were identified using CHOPCHOP (Labun et al., 2016; Labun et al., 2019). Oligonucleotides were annealed to each other and cloned into a pT7-gRNA plasmid, Addgene #46759 (Jao et al., 2013). Oligonucleotides utilized are shown in **Table S1**. gRNAs were synthesized from plasmids using a MEGAshortscript T7 Transcription Kit (Invitrogen AM1354) and purified.

One-cell-stage embryos were injected using glass needles pulled on a Sutter Instruments Fleming/Brown Micropipette Puller, model P-97 and a regulated air-pressure micro-injector (Harvard Apparatus, NY, PL1–90). Each embryo was injected with a 1nL solution containing one gRNA for *arnt1* and one gRNA for *arnt2* (30 ng/μL per target), Cas9 mRNA (150 ng/μL), and 0.1% phenol red. Mixtures were injected into the yolk of each embryo. Approximately 100 injected embryos per gRNA pair were raised to adulthood and crossed to AB zebrafish to generate F1 embryos. F1 offspring with heritable mutations were sequenced to identify mutations predicted to cause loss of function.

### Live Imaging

Embryos were imaged with a Nikon SMZ25 microscope equipped with a Hamamatsu ORCA-Flash4.0 digital CMOS camera, or with a Nikon SMZ18 microscope equipped with a Nikon DS-Fi3 camera. Images were equally adjusted for brightness and contrast in Adobe Photoshop CC 2020. Embryos and larvae were anesthetized with 0.04% tricaine and imaged in Petri dishes containing E3B. All embryos were genotyped after imaging.

### Whole-mount *in situ* hybridization (WISH)

We used the following DNA plasmids to generate RNA probes: *egfl7, lmo2* (Sun et al., 2018), *etsrp* (gift of Dr. Nathan Lawson, Addgene plasmid # 49005) (Moore et al., 2013)*, tal1* (gift of Dr. Arne Lekven)*, npas4l* (gift of Dr. Leonard Zon) (Reischauer et al., 2016)*, runx1* and *kdrl* (gifts of Dr. Mary Goll) (Li et al., 2015). For *lmo2* and *egfl7,* RNA was extracted from 10 pooled 1 dpf wild-type embryos using TRIzol Reagent (ThermoFisher Scientific, 15596026) and purified using RNA clean & concentration kit (Zymo Research, R1019). cDNA was synthesized using iScript cDNA synthesis (BioRad, 1708890). *lmo2* and *egfl7* were PCR amplified and Topo TA cloned into pCRII vector (Invitrogen, K4600-01). Primers used to amplify *lmo2* and *egfl7* can be found in **Table S1**. All DNA clones were verified by sequencing.

Digoxigenin-labeled antisense RNA probes were synthesized using T7 or SP6. Colorimetric *in situ* hybridization (ISH) was performed on fixed zebrafish embryos and larvae as previously described, with 5% dextran in the hybridization buffer (Lauter et al., 2011; Thisse and Thisse, 2008). Following ISH, embryos were washed several times in 1x PBST (phosphate buffered saline, 0.01% Tween), cleared with glycerol, or mounted in 3% methylcellulose on a class coverslip, and imaged on either a Nikon SMZ18 stereoscopic microscope or an inverted Nikon ECLIPSE Ti2-E microscope. Embryos were then genotyped. If embryos were older than 24 hpf, genomic DNA was extracted in 20 μL ELB (10 mM Tris pH 8.3, 50 mM KCl, 0.3% Tween 20) with 0.2 μL proteinase K (800 U/ml, NEB) in 96 well plates, one sample per well, at 55°C for 2 hours. If the embryos were 24 hpf or younger, they were first de- crosslinked in 300 mM NaCl for 4 hours at 65°C and held at 4-8°C until DNA could be extracted.

Genomic DNA was extracted using the HotSHOT protocol (Dobrzycki et al., 2018). Briefly, embryos were suspended in 10-20 μL of lysis buffer (50 mM NaOH) and incubated at 95°C for 30 minutes, cooled to 4°C, then 1/10^th^ the volume of neutralization buffer (1M Tris-HCl, pH 8.0) was added. Embryo lysate was incubated at 4°C overnight before starting PCR reactions. DNA regions of interest were amplified using one of two methods. The genes of interest were PCR amplified using JumpStart REDTaq ReadyMix (Sigma-Milipore, P0982) according to the manufacturer’s protocols. Alternatively, the DNA regions of interest were PCR amplified in two successive reactions using HS Taq (Takara, R007). The first PCR amplified a larger fragment of *arnt1* (about 700 bp) or *arnt2* (about 970 bp). Then the product of the larger fragment was used as a template to amplify a smaller fragment of *arnt1* (about 400 bp) or *arnt2* (about 650 bp). After PCR amplification, the PCR products were sequenced to determine the genotype. All primer sequences can be found in **Table S1**.

### Whole-mount *in situ* hybridization chain reaction (WIHCR)

HCR Antisense RNA probe sets for *arnt1, arnt2,* and *npas4l,* HCR amplifiers, and HCR buffers were obtained from Molecular Instruments, Inc (https://www.molecularinstruments.com/). Each gene had a probe set size of 20. WIHCR was performed on fixed AB embryos between 4-6 somites, as previously described (Choi et al., 2018; Ibarra-García-Padilla et al., 2021). Following HCR, embryos were washed several times in 1x PBST and cleared in glycerol. All embryos were mounted in 75% glycerol and imaged on a Nikon Ti2-E inverted microscope with Yokogawa CSU-W1 spinning disk confocal and Photometrics Prime95b sCMOS camera.

### WIHCR Image Processing and Analysis

All images used in processing and analysis were captured on the same day, using identical acquisition controls. We selected only images where a clear, bilateral view of the presumptive lateral plate mesoderm (LPM) and presumptive notochord was available (N=6 embryos from a single clutch, and N=4 negative control embryos from the same clutch). Images of anterior and posterior views of the embryos were collected. To calculate the number of 3D objects in a stack, we used Image J, FIJI version 2.3.0/1.53q (Schindelin et al., 2012), and the 3D-OC plug-in (Bolte and Cordelières, 2006). A custom threshold for each channel was applied to each image to filter out background autofluorescence seen in the negative control embryos. The number of detected objects for probed samples was then normalized to negative control by subtracting the average number of detected objects from control embryos from the number of detected objects in probed embryos. We used ImageJ to calculate the adjusted integrated intensity of probed and control embryos (Schindelin et al., 2012). First, maximum intensity projections (MIP) of the z-stacks were created, then the region of interest (ROI) was drawn around the areas with the highest concentration of *npas4l* expression (presumptive LPM) or around the corresponding region of the presumptive LPM in the negative control embryos. The percent area of pixels and the mean gray value for each channel were calculated and limited to the threshold. The adjusted integrated density was then calculated by multiplying the percent area and the mean gray value.

### O-dianisidine labeling of blood cells

Zebrafish embryos were stained at approximately 72 hours post fertilization. Embryos were dechorionated using 0.2% pronase in E3B media for 5-10 minutes and were gently triturated with a 1000µl pipette. Embryos were transferred to glass vials and submerged in o-dianisidine working solution, prepared as described (Kafina and Paw, 2018). Vials were kept in the dark for 5 minutes during staining. Samples were then fixed in 4% formaldehyde for 30 minutes at room temperature. Embryos were washed in PBS-Tween 0.1% before mounting for imaging in 3% methylcellulose on a glass dish or coverslip. Embryos were genotyped after imaging.

### Plasmid synthesis for microinjection and *in vitro* transfection

Cloning of plasmids for expression in cultured cells was performed as follows. For npas4l-V5, we took pCS2+npas4lORF (gift of Dr. Leonard Zon) that contains the open reading frame (ORF) of zebrafish *npas4l* downstream of the CMV promoter. This plasmid was modified to include a C-terminal V5 tag using Q5 Site-Directed Mutagenesis Kit (New England Biolabs, #E0554S). For arnt1-myc, we commercially synthesized a plasmid containing CMV promoter driving a codon-optimized zebrafish *arnt1* ORF (ENSDART00000081852.5) with a C-terminal myc tag (Twist Bioscience, San Francisco, CA). For arnt2-myc, we commercially synthesized a gene block of the zebrafish *arnt2* ORF (ENSDART00000158162.2, Integrated DNA Technologies). The gene block was integrated into a plasmid containing a CMV promoter using pcDNA3.3-TOPO TA cloning kit (Invitrogen, K8300-01) in accordance with manufacturer’s instructions. A C-terminal HA tag was added using Q5 Site-Directed Mutagenesis Kit (New England Biolabs, #E0554S). For microinjection, an SP6 promoter was inserted upstream of the transcriptional start using Q5 Site-Directed Mutagenesis Kit (New England Biolabs, #E0554S). All plasmids were confirmed by sequencing.

### Cell Culture and Transient Transfection

HEK293T cells (gift from Dr. Charles Foulds), were thawed and maintained at 37°C in humidified 5% CO2 atmosphere incubator. Cells were cultured in DMEM supplemented with 10% fetal bovine serum (FBS; Gibco, 2062564) and 1% penicillin-streptomycin (Gibco, 15140-122). After 24 hours of passaging in a 100mm plate at a density of 2.2 x 10^6^ cells per dish and reaching at least 70 confluency, the media was changed to DMEM supplemented with 1% dialyzed FBS. Cells were transiently transfected with 12 μg of total DNA and Lipofectamine 2000 according to the manufacturer’s instructions (Invitrogen, 2400439) with *npas4l-v5* together with either *arnt1-myc* or *arnt2-HA* plasmid DNA.

### Co-immunoprecipitation and Western blot

Transiently transfected cells were washed with ice cold PBS and lysed in 500 μL of 1x RIPA buffer containing Pierce Protease (ThermoScientific, A32955). Cells were sonicated in 6 x 30 second intervals using a Bioruptor 300 Machine (Diagenode), then centrifuged for 10 minutes at 10,000g at 4°C. The lysate (supernatant) was removed from the pellet and placed in fresh 1.5mL Eppendorf tube. Cell lysate was precleared with Protein A beads (Invitrogen #159 18-014) for 30 minutes 4°C, incubated with either 5 μg of V5 antibody (ThermoFisher Scientific R960-25) or myc antibody (SantaCruz, SC-40) diluted 1:200 for 1 hour, then immunoprecipitated with 20 μL of Protein A beads for 1 hour at 4°C. The beads were washed three times with RIPA buffer and once with PBS and finally suspended in 30μL of PBS. Equal volume of 2x Laemmli Sample Buffer (BioRad, cat# 1610737) was added to each sample and boiled for 5 minutes at 95°C then placed on ice. Samples were run on 4-20% Mini-PROTEAN TGX gels (BioRad, cat# 4561093) and transferred onto PVDF membrane using Trans-Blot Turbo Transfer System (BioRad) using the pre-defined Mixed MW program. The membranes were washed several times with 1x Tris-Buffered Saline, 0.1% Tween 20 (TBST) (GBiosciences, R043) and incubated for 1 hour with agitation in 5% milk in 1x TBST. Membranes were probed with 1:1000 V5-HRP (ThermoFisher Scientific, R960-25), 1:1000 myc-HRP (SantaCruz, SC-40), or 1:200 HA (Cell Signaling, 3724) antibodies rocking overnight at 4°C. Membranes were washed 5 times with 1xTBST on orbital rocker for 15 minutes at room temperature. Membranes probed with the anti-HA antibody were then probed with 1:10,000 IRDye 800CW donkey anti-rabbit (Licor, 926-32213) in 2.5%-5% milk in 1x TBST for 1 hour and then washed multiple times with 1xTBST on orbital rocker for 5 minutes prior to imaging. Membranes probed with HRP conjugated antibodies were developed using Clarity™ Western ECL Substrate (BioRad, 170-5061). Membranes were imaged using BioRad ChemiDoc™ MP Imaging System.

### Zebrafish embryo injection

Embryo injections were performed as described (Westerfield, 2000) with the following modifications. Plasmids containing either *npas4l, arnt1,* or *arnt2* were linearized by restriction enzyme digestion. mRNA was transcribed using the mMessage mMachine kit (Invitrogen, AM1340) and purified with the RNA clean & concentration kit (Zymo Research, R1019). Embryos were microinjected at the 1-cell stage with 1 nL total volume containing either *npas4l* mRNA (50 pg mRNA) or a mixture of *arnt1* and *arnt2* mRNA (50 pg each for 100 pg total mRNA). All solutions were injected into the yolk.

### TCDD exposure

At 1-day post fertilization (dpf), embryos were exposed to 10 ng/mL 2,3,7,8-tetrachlorodibenzodioxin (TCDD, AccuStandard Inc, D404N) or vehicle (0.1% DMSO). All treatments were performed on a single clutch of at least 20 embryos. Embryos treated with vehicle were from the same clutches as embryos treated with TCDD. Chemical exposures were performed on at least 2 clutches per genotype. Embryos were raised at 28.5°C on a 14-h light, 10-h dark cycle until 3 dpf, when they were imaged and then genotyped.

### Statistical Analysis

All Statistical analyses were performed using Prism version 9.4.1 (GraphPad Software, Boston, MA). Mendelian Frequencies Analysis: For the Chi-squared analysis of embryo and adult genotype ratios we eliminated all 0 values from the analysis.

#### WISH Phenotype Frequency Analysis

For WISH images of *arnt1/2* mutants, we used a binomial test to compare the observed distribution of embryos with the reduced expression phenotype to the calculated expected number of *arnt1/2* mutants. Additionally, we performed a Spearman rank correlation analysis on genotyped embryos to determine if there was a correlation between the number of mutant alleles and the observed phenotype. Each genotype was assigned a number 0-4: with 0 indicating no mutant alleles (wild-type siblings), 1 indicating one mutant allele (*arnt1* heterozygous and *arnt2* heterozygous siblings), 2 indicating two mutant alleles (*arnt1* homozygous, *arnt2* homozygous, or *arnt1/2* heterozygous siblings), 3 indicating three mutant alleles (*arnt1* homozygous; *arnt2* heterozygous and *arnt1* heterozygous; *arnt2* homozygous siblings), and 4 indicating 4 mutant alleles (*arnt1/2* mutants). The overserved phenotypes were assigned a number value: normal expression was coded as a 1 and reduced expression was coded as 0.

#### WIHCR Image Analysis

For WIHCR image analysis, the variance was not assumed for *npas4l* and *arnt2* samples. *arnt1* samples did not have a normal distribution, thus a non-parametric Mann-Whitney test was performed. We were able to identify an outlier in the *arnt1* integrated intensity data using Grubb’s test (α=0.05), however, when this outlier was removed from analysis there was no change in significance or Mann-Whitney U value.

#### TCDD Treatment Analysis

For TCDD treated embryos, unpaired Welch’s *t-*tests were performed to compare the average number of embryos with cardiotoxicity to the average number of embryos with a normal phenotype from a given genotype.

## RESULTS

### *arnt1/2* homozygous mutants lack circulating blood

To determine if *arnt1* or *arnt2* is required for hemato-vascular development, we generated heritable mutant alleles for *arnt1 (arnt1^bcm1^* and *arnt1^bcm2^)* and *arnt2 (arnt2^bcm3^)* using CRISPR/Cas9. For *arnt1^bcm1^,* we generated zebrafish with a 5 bp insertion in exon 5, which results in a frameshift at amino acid (AA) 126 and a premature stop codon in the first PAS domain at AA139. For *arnt1^bcm2^*, we generated zebrafish with an 11 bp deletion in exon 10, which results in a frameshift at AA230 and a premature stop codon between the two PAS domains at AA250 (**Figure 1B**). We did not observe any phenotypic differences between homozygous *arnt1^bcm/bcm1^* and *arnt1^bcm2/bcm2^*mutant embryos or larvae, these fish will herein be called *arnt1* mutants. During embryogenesis and larval development, *arnt1* mutants appear to have a normal phenotype and are indistinguishable from age-matched wild-type larvae (**Figure 1D-E**). *arnt1* mutants are viable to adulthood and are fertile, consistent with previously published *arnt1* mutants (Marchi et al., 2020). Maternal zygotic *arnt1* mutants did not show a developmental phenotype and were indistinguishable from wild-type or zygotic *arnt1* mutants. For *arnt2^bcm3^,* we generated zebrafish with a 3 bp deletion and 1 bp insertion in exon 9, which resulted in a missense mutation at AA286 and a premature stop codon between the two PAS domains at AA287 (**Figure 1C**). During embryogenesis and larval development, *arnt2^bcm3/bcm3^* embryos, herein called *arnt2* mutants, were indistinguishable from wild-type embryos (**Figure 1D-F**). *arnt2* mutants were viable at 5 days post fertilization (dpf) but did not survive to adulthood, consistent with other independently generated *arnt2* mutant zebrafish lines (Hill et al., 2009; Löhr et al., 2009).

Because Arnt1 and Arnt2 can interact with multiple Class I bHLH-PAS proteins (**Figure 1A**), we examined if embryos that lacked both *arnt1* and *arnt2* had a hemato-vascular phenotype. To generate these embryos, we crossed *arnt1^bcm2/+^;arnt2^bcm3/+^*adult fish to each other. We observed all genotypes at the expected Mendelian ratios in offspring at 24 to 48 hours post fertilization (**Table 1**). However, *arnt1^bcm2/bcm2^; arnt2^bcm3/bcm3^* embryos, herein called *arnt1/2* mutants, showed a clear phenotype (**Figure 1G**). These embryos lack all circulating blood (**Figure 1H-L, and Movies 3, 4, 7, 8**). All other intermediate *arnt1;arnt2* genotypes, e.g., *arnt1^bcm2/+^;arnt2^bcm3/+^*, appeared phenotypically wild-type (**Figure 1H, J, K, and Movies 1, 2, 5, 6**). These results suggest that *arnt1* and *arnt2* function redundantly because only the *arnt1/2* double homozygous mutants show a phenotype. Additionally, the *arnt1/2* mutant phenotype appears identical to the previously described *npas4l* mutant phenotype, also known as *cloche* (Reischauer et al., 2016; Stainier et al., 1995). Like *npas4l* mutants, *arnt1/2* mutants exhibit pericardial edema and have an enlarged heart atrium, likely a secondary effect of the lack of blood and endocardial cells (**Figure 1G, Movies 3, 7**). Taken together, these data suggest that *arnt1* and *arnt2* have a similar function as *npas4l*.

Next, we examined the frequencies of the genotypes in adult fish from *arnt1^bcm2/+^; arnt2^bcm3/+^* parents. Given that *arnt2* mutants are not viable as adults, these adults can be one of six possible genotypes. Interestingly, we did not observe the expected Mendelian ratios of alleles in these adult fish (**Table 2**; *X^2^*(df=5, N=138) = 45.57, p<0.0001, N=138 offspring from two different clutches). There is a reduction in the number of observed *arnt1^bcm2/bcm2^*;*arnt2^bcm3/+^* heterozygous adults compared to the expected number. We hypothesize that the number of *arnt1* and *arnt2* wild-type alleles correlates with the likelihood of juvenile fish surviving to adulthood.

**Table 2.** Genotype frequencies of viable adult fish from arnt1^bcm2/+^; arnt2^bcm3/+^ parents. Offspring were raised to adulthood (≥3 months old) and genotyped. The expected Mendelian ratios were adjusted to reflect the fact that arnt2^bcm3/bcm3^ mutants are not viable to adulthood. The frequencies of the observed genotypes were significantly different from the expected genotypes, with fewer arnt1^bcm2/bcm2^; arnt2^bcm3/+^ observed. X^2^ (5, N=138) = 45.57, p<0.0001.

| Genotype | Expected Percentage | Expected n=138 | Actual Percentage | Actual n=138 |
| --- | --- | --- | --- | --- |
| *arnt1*<sup>+/+</sup>; *arnt2*<sup>+/+</sup> | 7.69% | 10.6 | 7.24% | 10 |
| *arnt1*<sup>+/+</sup>; *arnt2*<sup>bcm3/+</sup> | 15.4% | 21.25 | 15.2% | 21 |
| *arnt1*<sup>+/+</sup>; *arnt2*<sup>bcm3/bcm3</sup> | 0% | 0 | 0% | 0 |
| *arnt1*<sup>bcm2/+</sup>; *arnt2*<sup>+/+</sup> | 13.3% | 18.4 | 30.4% | 42 |
| *arnt1*<sup>bcm2/+</sup>; *arnt2*<sup>bcm3/+</sup> | 30.8% | 42 | 43.5% | 60 |
| *arnt1*<sup>bcm2/+</sup>; *arnt2*<sup>bcm3/bcm3</sup> | 0% | 0 | 0% | 0 |
| *arnt1*<sup>bcm2/bcm2</sup>; *arnt2*<sup>+/+</sup> | 7.69% | 10.6 | 1.45% | 2 |
| *arnt1*<sup>bcm2/bcm2</sup>; *arnt2*<sup>bcm3/+</sup> | 15.4% | 21.25 | 2.17% | 3 |
| *arnt1*<sup>bcm2/bcm2</sup>; *arnt2*<sup>bcm3/bcm3</sup> | 0% | 0 | 0% | 0 |

### *arnt1/2* mutants have reduced expression of hematopoietic and endothelial cells

We hypothesized that the Arnt proteins are dimerization partners with Npas4l and that *arnt1/2* mutants would exhibit the same phenotypes as *npas4l* mutants. Therefore, we examined *arnt1/2* mutant embryos for additional phenotypes found in *npas4l* mutants. Previous studies determined that *npas4l* mutants lack almost all endothelial cells, except a population of endothelial cells located in the lower trunk and caudal tail of the larvae (Liao et al., 1997; Reischauer et al., 2016; Stainier et al., 1995). To determine if our *arnt1/2* mutants showed a similar reduction in endothelial cells, we performed whole-mount *in situ* hybridization (WISH) on 2 dpf embryos for *kdrl* (*flk-1)*. *kdrl*, a widely used marker of endothelial cells in blood vessels, is essential for the development of blood vessels in mice and zebrafish (Dumont et al., 1995; Liao et al., 1997; Shalaby et al., 1997). We found that *arnt1* mutants, *arnt2* mutants, and wild-type embryos showed clear expression of *kdrl* in blood vessels at 2 dpf (**Figure 2A-C**, N=2-5 biological replicates per genotype per probe, 10-100 embryos examined per biological replicate). In contrast, *arnt1/2* mutants had a reduced number of *kdrl-*positive cells at 2 dpf (**Figure 2D**). The *kdrl*-positive cells present in the *arnt1/2* mutant were in the caudal tail and likely comprised a portion of the caudal vein (**Figure 2D’**). To determine if this population of cells was present earlier in development we bred *arnt1^bcm2/+^; arnt2^bcm3/+^* onto a Tg(*fli1:egfp^y1^)* transgenic background, which labels all blood vessels and the pharyngeal arches of the developing embryos. At 1 dpf, *arnt1* mutants, *arnt2* mutants, and wild-type embryos had GPF-positive vasculature structures (**Figure 2I-K**, N=2 biological replicates, with 21-131 embryos per clutch, 232 embryos total). Within the tails of these embryos the dorsal aorta, caudal vein, and intersegmental vessels are clear and present (**Figure 2I’-K’**). In contrast, *arnt1/2* mutants had reduced GFP-positive vascular structures (**Figure 2L).** Within these embryos, only the extreme caudal vein was fluorescent (**Figure 2L’**). This population of endothelial cells is also seen in *npas4l* mutants (Liao et al., 1997). Together these results suggest that *arnt1* or *arnt2* is required for *npas4l-*dependent blood vessel development.

**Figure 2.**
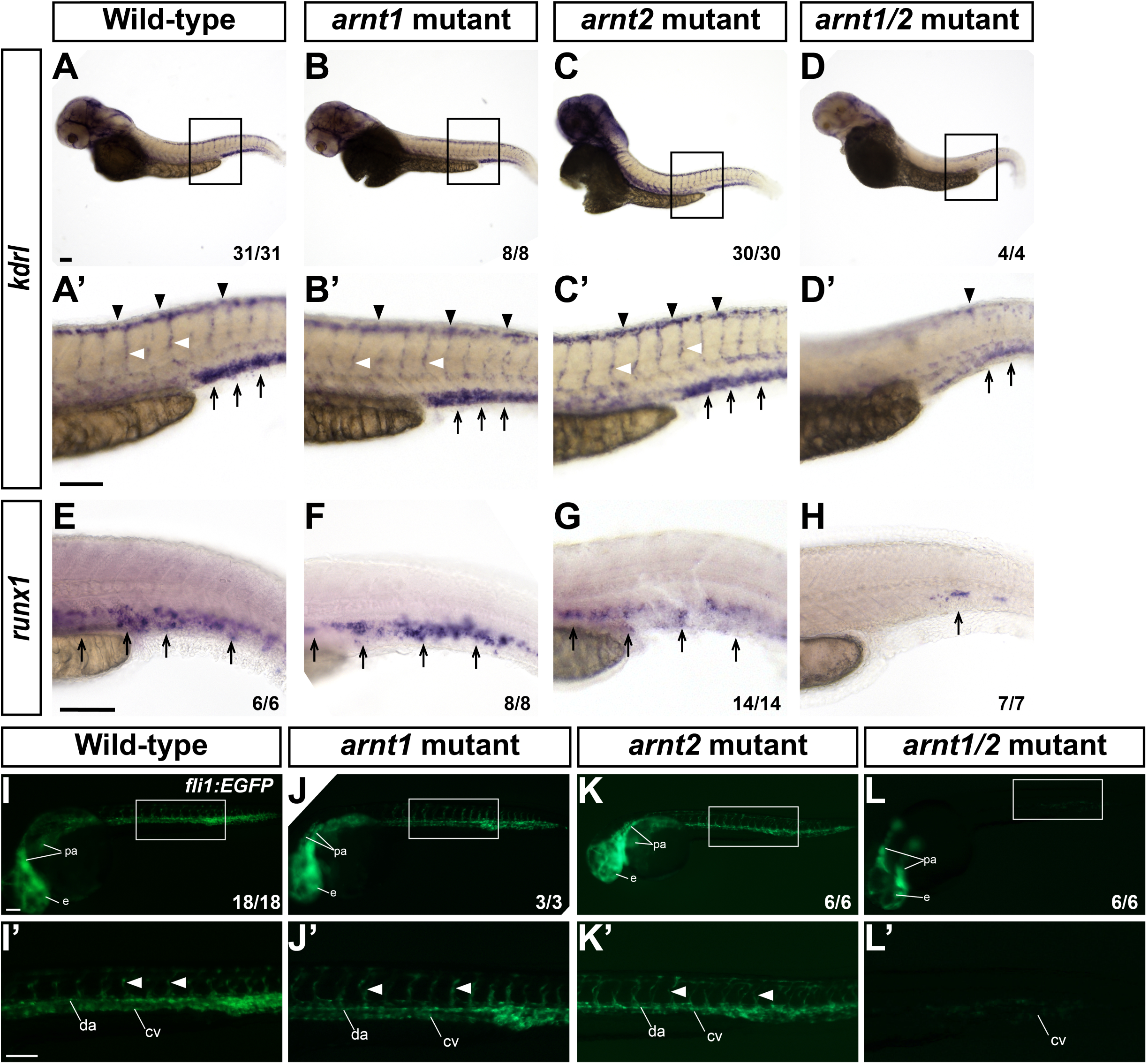
*arnt1/2* mutants show reduced levels of endothelial cells and *runx1*-positive hematopoietic cells. (A-L) Embryos were probed for *kdrl* (A-D) or *runx1* (E-H) using whole-mount in situ hybridization at 2 days post fertilization (dpf). (A-D) *kdrl* staining in wild-type, *arnt1* mutant, *arnt2* mutant, and *arnt1/2* mutant embryos. (A’-D’) Images of each embryo’s trunk near the end of the yolk extension (indicated by the box in panels A-D). Black arrowheads indicate *kdrl* staining in the dorsal longitudinal anastomotic vessels, black arrows indicate the dorsal aorta and caudal vein, and white arrowheads indicate inter-segmental vessels. (E-H) Black arrows indicate *runx1* labeling in the presumptive hematopoietic stem cell niche. (I-L) Wild-type and mutant embryos on Tg(*fli1:EGFP ^y1^)* transgenic background were imaged at 1 dpf and then genotyped. (I’-L’) Enlarged images of each embryo’s trunk near the end of the yolk extension (indicated by the box in panels I-L). White arrowheads correspond to inter-segmental vessels. e: eye, pa: pharyngeal arch, da: dorsal aorta, cv: caudal vein. The fraction in the bottom right corner indicates the number of embryos of the given genotype with the represented phenotype over the total number of embryos of the given genotype from a single clutch. Scale bars,100 μm.

In addition to a clear reduction of endothelial cells, *npas4l* mutants show a reduction in the number of hematopoietic stem cells (HSC) (Stainier et al., 1995; Thompson et al., 1998). To determine if *arnt1/2* mutants contain HSCs, we examined *runx1* expression by WISH. *runx1* is a marker of HSCs and is required for definitive hematopoiesis in zebrafish (Kalev-Zylinska et al., 2002). At 48 hpf, *runx1-* positive HSCs are known to localize to the area between the dorsal aorta and the posterior caudal vein (Lam et al., 2009; North et al., 2009). At 2 dpf, *arnt1* mutants, *arnt2* mutants, and wild-type embryos all contain *runx1-*positive cells near the dorsal aorta (**Figure 2E-G**). However, *arnt1/2* mutants have little or no *runx1* labeling in this region (**Figure 2H**). This suggests that *arnt1* or *arnt2* is necessary for hematopoietic cell development during embryogenesis, just like *npas4l* (Liao et al., 1997; Liao et al., 1998; Stainier et al., 1995; Thompson et al., 1998).

### Hematopoietic and endothelial progenitor populations are reduced in *arnt1/2* mutants

Hemato-vascular progenitor specification from the multipotent mesoderm begins in mid- gastrula, around shield stage, and peaks during segmentation (Vogeli et al., 2006). Npas4l is thought to directly upregulate target genes that cause cells to differentiate into endothelial cells and hematopoietic stem and progenitor cells. If Npas4l, Arnt1 and Arnt2 act in tandem to regulate target gene expression, then we expect that Npas4l-target genes would be downregulated in *arnt1/*2 mutants. Therefore, we hypothesized that Npas4l-target genes would be downregulated in *arnt1/*2 mutants. We focused on four putative *npas4l* target genes known to be important for the hemato-vascular lineage: *etsrp (etv2), tal1, lmo2 (scl),* and *egfl7* (Marass et al., 2019; Sumanas and Lin, 2006; Sumanas et al., 2005). We bred *arnt1^bcm2/+^; arnt2^bcm3/+^* adults to each other to generate embryos that were *arnt1/2* mutant, *arnt1* mutant, *arnt2* mutant, wild-type, and all other intermediate genotypes (referred to as *arnt1/2* siblings). At the 2-6 somite stages (about 10-14 hpf), *npas4l* mutant embryos show a reduction of expression in *etsrp*, *tal1,* and *lmo2* (Reischauer et al., 2016; Thompson et al., 1998). We examined the expression of these genes in *arnt1/2* mutants and *arnt1/2* siblings between 4-6 somite stage (**Figure 3**). We found that 7% of embryos from *arnt1^bcm2/+^;arnt2^bcm3/+^* parents showed reduced expression of *etsrp* compared to siblings, (**Figure 3A-B**, *N=*4 biological replicates, 80-250 embryos per clutch, 604 embryos total). This did not differ from the expected Mendelian ratio of *arnt1/2* mutants (6.25%; binomial test, *p>*0.05). To confirm that these embryos were in fact *arnt1/2* mutants, following WISH we genotyped 24 embryos with reduced expression of *etsrp* and found that 100% were *arnt1/2* mutants. We also genotyped 26 embryos with normal expression of *etsrp,* only two of which were *arnt1/2* mutants (**Table S2**). To examine if there is an association between the number of mutant alleles and the observed phenotype, we performed a Spearman Rank correlation test (see Methods for details) and found that there is a statistically significant correlation between the number of mutant alleles and the observed phenotype (*Spearman correlation,* r= -0.9, *p*<0.001). We conclude that the expression of *etsrp* is reduced in *arnt1/2* mutants compared to wild-type.

**Figure 3.**
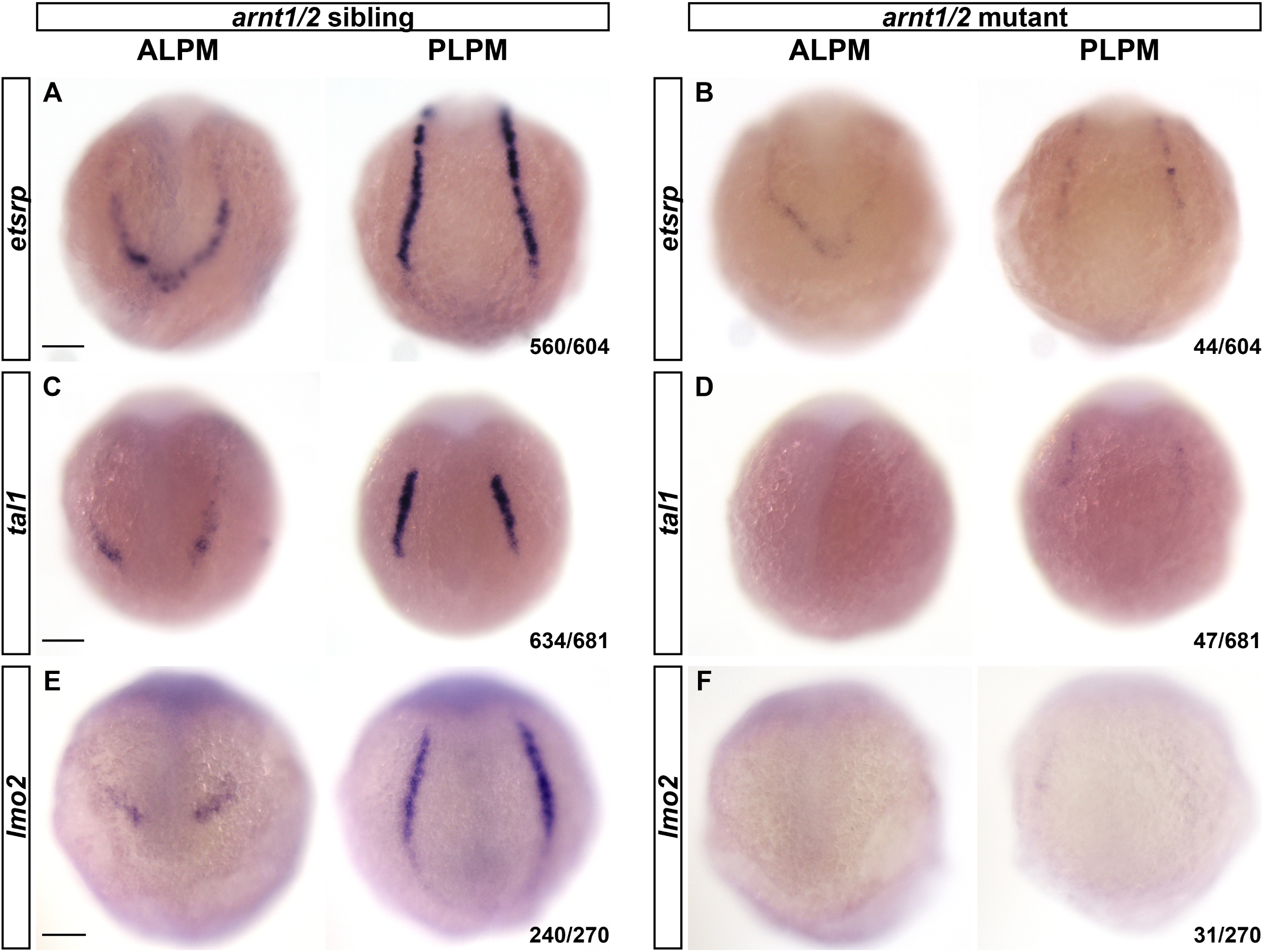
*arnt1/2* mutants show decreased *etsrp*, *tal1*, and *lmo2* expression in the lateral plate mesoderm at 4-6 somite stages. Whole-mount in situ hybridization for *etsrp*, *tal1,* and *lmo2* expression was performed on 4-6 somite stage embryos that were offspring of *arnt1^bcm2/+^;arnt2^bcm3/+^* adult zebrafish. (**A**) Expression of *etsrp* was observed in the anterior and posterior lateral plate mesoderm (ALPM, PLPM) in 93% of embryos. (**B**) Limited or no expression of *etsrp* was observed in 7% of embryos, consistent with the expected Mendelian ratio of 6.25% for *arnt1^bcm2/+^;arnt2^bcm3/+^* (*arnt1/2* mutants). (**C**) Expression of *tal1* was observed in the ALPM and PLPM in 93% of embryos. (**D**) Limited or no expression of *tal1* was observed in 7% of embryos, consistent with the expected Mendelian ratio of 6.25% for *arnt1/2* mutants. (**E**) Expression of *lmo2* was observed in 89% of embryos. (**F**) Limited or no expression of *lmo2* was observed in 11% of embryos, which is slightly more than the expected Mendelian ratio of 6.25% for *arnt1/2* mutants. We genotyped a subset of each group of embryos (see **Supplemental Tables 2-4**) and observed a correlation between number of mutant alleles and expression of *etsrp*, *tal1*, and *lmo2* genes. Spearman correlation, *etsrp* r= -0.9, p<0.001; *tal1* r= -0.6, p<0.005; *lmo2* r= -0.2, p<0.01. All embryos are oriented with dorsal towards the top. Fractions in bottom right corners refer to the number of embryos with the indicated phenotype over the total number of embryos examined. Scale bars, 100 μm

We found that 7% of embryos from arnt*1^bcm2/+^; arnt2^bcm3/+^* parents showed reduced expression of *tal1* compared to *arnt1/2* siblings (**Figure 3C-D**, N=6 biological replicates, 60-300 embryos per clutch, 681 embryos total). This did not differ significantly from the expected Mendelian ratio of *arnt1/2* mutants (binomial test, *p>*0.05). To confirm these embryos were in fact *arnt1/2* mutants, we genotyped 18 embryos with reduced expression and confirmed that 83% (15 of 18 embryos) of these embryos were *arnt1/2* mutants (**Table S3**). We genotyped 11 embryos with normal expression of *tal1,* none of which were *arnt1/2* mutants (**Table S3**). To examine if there is a relationship between the number of mutant alleles and the observed phenotype, we performed a Spearman Rank correlation test and found that there is a statistically significant correlation between the number of mutant alleles and the observed phenotype (*Spearman correlation,* r= -0.6, *p*<0.005). We conclude that the expression of *tal1* is reduced in *arnt1/2* mutants compared to wild-type.

We found that 11% of embryos from *arnt1^bcm2/+^;arnt2^bcm3/+^*parents showed reduced expression of *lmo2* compared to siblings (**Figure 3D-E**, N*=*2 biological replicates, 95-175 embryos per clutch, 270 embryos total). This did significantly differ from the expected 6.25% Mendelian ratio of *arnt1/2* mutants (binomial test, *p<*0.01). To test for a phenotype-genotype correlation, we genotyped 185 embryos following labeling for *lmo2:* 160 embryos with normal *lmo2* expression and 25 embryos with reduced *lmo2* expression. Of the 25 embryos with reduced expression of *lmo2*, we observed 8 embryos that were *arnt1/2* mutants, 8 embryos with three mutant alleles, 4 embryos with two mutant alleles, 3 embryos with one mutant allele, and 2 wild-type embryos (**Table S4**). Of the 160 embryos with normal expression of *lmo2*, 5 embryos were *arnt1/2* mutants, 36 embryos had three mutant alleles, 64 embryos had two mutant alleles, 46 embryos had one mutant allele, and 9 were wild-type embryos (**Table S4**). To examine if there is a relationship between the number of mutant alleles and the observed phenotype, we performed a Spearman rank correlation and found that there is a statistically significant correlation between the number of mutant alleles and the observed phenotype (*Spearman correlation*, r = -0.2, p<0.01). We conclude that the expression of *lmo2* is reduced in *arnt1/2* mutants compared to wild-type.

We examined the expression of *egfl7* at 10-somite stage embryos from *arnt1^bcm2/+^; arnt2^bcm3/+^* parents. We observed *egfl7* expression in portions of the ALPM and PLPM in wild-type and in 95% of the embryos from *arnt1^bcm2/+^; arnt2^bcm3/+^* parents, but no expression of *egfl7* in the anterior of 5% of embryos from *arnt1^bcm2/+^; arnt2^bcm3/+^* parents, N=11 of 214 embryos (**Figure S1**, N=2 biological replicates, 94-120 embryos per clutch, 214 total embryos, and age-matched wild-type embryos, N=1 biological replicate, 28 embryos). This did not differ from the expected Mendelian ratio of *arnt1/2* mutants (6.25%; binomial test, *p>*0.05). To examine if there is a relationship between the number of mutant alleles and the observed phenotype, we performed a Spearman Rank correlation test and found that there is a statistically significant correlation between the number of mutant alleles and the observed phenotype (*Spearman correlation,* r= -0.5, *p*<0.001) (**Table S5**). We conclude that the expression of *egfl7* is reduced in *arnt1/2* mutants compared to wild-type.

These results support the idea that *npas4l*, *arnt1* and *arnt2* regulate identical target genes for hemato-vascular specification. *arnt1* and *arnt2* could interact with *npas4l* at the same level of the pathway, consistent with bHLH-PAS transcription factors acting as heterodimers. Alternatively, it is possible that *arnt1* and *arnt2* act upstream of *npas4l*. To distinguish among these ideas, we tested whether *npas4l* expression was reduced in *arnt1/2* mutants. At the 2-4 somite stage, *arnt1/2* mutants exhibit normal expression of *npas4l* in the lateral plate mesoderm, indistinguishable from age-matched wild-type embryos (**Figure S2**, **Table S6**, N=2 biological replicates, 68-100 embryos per clutch, 168 total embryos). This result indicates that *npas4l* expression occurs independently of *arnt1* and *arnt2*. We conclude that *npas4l* does not act downstream of *arnt1* or *arnt2*. Taken together, our results suggest that *arnt1* and *arnt2* act in conjunction with *npas4l* during hemato-vascular specification.

### *arnt1* and *arnt2* expression is ubiquitous and colocalizes with *npas4l* expression

If *arnt1, arnt2* and *npas4l* act together to specify hemato-vascular progenitor cells, then all three genes should be expressed in the same cells in the lateral plate mesoderm. To determine if *arnt1, arnt2,* and *npas4l* are expressed in the same cells, we performed whole-mount *in situ* hybridization chain reaction (WIHCR) on wild-type zebrafish embryos at the 4-somite stage (approximately 10.5 hpf). At this stage, *npas4l* is robustly expressed in the LPM (Reischauer et al., 2016). We found that *arnt1* and *arnt2* are expressed ubiquitously throughout the embryo at this stage, although both genes appear to be expressed at low levels (**Figure 4A and Figure S3A**, N=3 biological replicates, 5-9 embryos per replicate, 22 embryos total).

**Figure 4.**
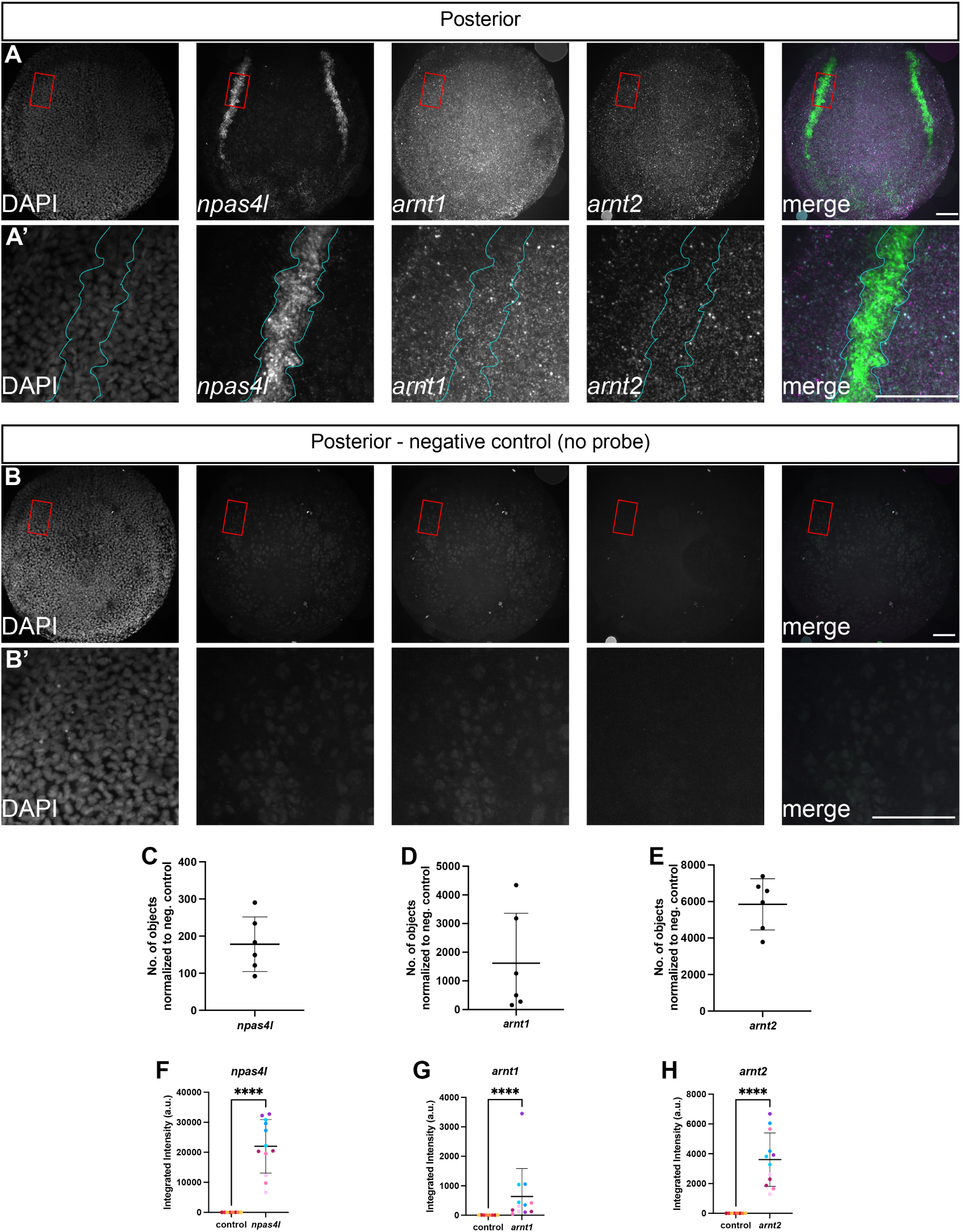
*arnt1* and *arnt2* are ubiquitously expressed at the 4-somite stage and co-localized with *npas4l* expression. Whole-mount *in situ* hybridization chain reaction was performed on wild-type (AB) embryos at the 4 somite stage. (**A**) Embryos were probed for *npas4l, arnt1,* and *arnt2* expression and counter-stained with DAPI. *arnt1* and *arnt2* labeling was detected ubiquitously, while *npas4l* labeling was localized to the lateral plate mesoderm. (A’) Higher magnification views of the boxed areas in (A) shows co-localization of *arnt1, arnt2,* and *npas4l* in the posterior lateral plate mesoderm, outlined in cyan. (**B**) To demonstrate that the observed expression of *arnt1* and *arnt2* is not autofluorescence of the embryo or the yolk, wild-type embryos were processed identically to those in (A), but no anti-sense RNA probe sets were added. (B’) Higher magnification views of the boxed areas in (B) shows what background/autofluorescence looks like in the presumptive posterior lateral plate mesoderm. All images in (**A, B**) are maximum intensity projections from z-stacks taken every 2.5 μm. In merged panels, *npas4l* is green, *arnt1* is magenta, *arnt2* is cyan. The posterior portion of the embryo is in view, dorsal is towards the top of the image. Scale bars, 100 μm. (**C-E**) To validate the observed expression of *arnt1* and *arnt2,* we quantified the number of detectable objects above a set threshold in the whole embryo across each z-plane for 4 negative control embryos and 6 experimental embryos probed for *arnt1*, *arnt2* and *npas4l*. We then normalized the number of detected objects by subtracting the average number of objects detected in the negative controls from the number of detected objects in the experimental embryos. Each point on these graphs represents the total number of objects above threshold for a single embryo. (**F-H**) To demonstrate that *arnt1* and *arnt2* expression occurs in the lateral plate mesoderm (LPM), we calculated the integrated intensity of fluorescence (pixel area, pixel intensity) over each side of the posterior LPM in maximum intensity projections from control and experimental embryos. Each point represents the integrated intensity from one side of the posterior LPM from a single embryo (right or left side), where each embryo has two bilateral *npas4l*-positive regions of the posterior LPM. Points of the same color are measurements from opposite sides of the posterior LPM from the same embryo. *npas4l*, *arnt1*, and *arnt2* labeling was statistically significantly increased in the lateral plate mesoderm compared to negative control embryos. *npas4l* Welch’s t-test, t(11)=8.6, p<0.0001; *arnt1* Mann-Whitney, U=0, p<0.0001; *arnt2* Welch’s t-test, t(11)=7.0, p<0.0001

Because no previous publications examined the spatial distribution of *arnt1* or *arnt2* mRNA in embryos younger than 24 hpf (Löhr et al., 2009), we tested whether background or autofluorescence influenced our results. Samples probed for *arnt1*, *arnt2,* and *npas4l* were compared to negative control embryos, embryos from the same clutch that underwent the same WIHCR protocol, but in the absence of anti-sense RNA probe sets (**Figure 4B and Figure S3B**, N=3 biological replicates, 6-12 embryos per replicate, 28 embryos total). We verified that the observed *npas4l*, *arnt1,* and *arnt2* expression was specific using two different image analysis tools. First, we analyzed representative images in 3- dimensions using the ImageJ plug-in “3D Count Objects” (Bolte and Cordelières, 2006). The number of objects above a set threshold was measured for each channel and then normalized to background by subtracting the average number of objects detected in the corresponding channel of the negative control embryos. *npas4l-, arnt1-*, and *arnt2-*positive objects were all detected in the posterior of the embryos (**Figure 4C-E**). Second, we analyzed the expression of *arnt1* and *arnt2* in the presumptive LPM by drawing regions of interest (ROIs) encompassing the observed *npas4l* signal, and then measured the percent area of expression and the mean gray intensity above the threshold for each channel. The integrated intensity was calculated by multiplying the percent area by the mean grey intensity, thus taking both brightness and area of signal into consideration (**Figure 4F-H, Figure S3 C- E**). We found that *npas4l, arnt1,* and *arnt2* all showed statistically significantly more expression in experimental embryos compared to the negative control, as measured by both number of objects and integrated intensity of objects in the LPM (**Figure 4F-H**, *npas4l* Welsh’s t-test, p<0.0001; *arnt2* Welsh’s t-test, p<0.0001; *arnt1* Mann-Whitney test, *p<0.0001*).

We asked whether the expression of *npas4l*, *arnt1* and *arnt2* was the same in the anterior LPM (ALPM) and posterior LPM (PLPM). *npas4l* and its associated downstream targets exhibit more expression in the PLPM compared to the ALPM in embryos throughout segmentation (Liao et al., 1997; Liao et al., 2000; Marass et al., 2019; Mattonet et al., 2022; Qian et al., 2005; Stainier et al., 1995). At 4- 6 somite stage embryos, we found that expression of *npas4l* and *arnt2* was greater in the PLPM than in the ALPM, but the expression of *arnt1* was similar in both regions (**Figure S3C, S3E,** *npas4l* Welsh’s t- test, p<0.0001; *arnt2* Welsh’s t-test, *p<0.005*). We conclude that *arnt1* and *arnt2* are expressed within the *npas4l-*positive LPM, consistent with our hypothesis that all 3 genes together regulate the specification of hemato-vascular progenitor cells.

### Expression of *npas4l*-target genes requires *npas4l*, *arnt1* and *arnt2*

*npas4l* is not expressed until tailbud stage (Reischauer et al., 2016). A previous study demonstrated that ectopic expression of *npas4l* in shield-stage embryos, prior to the onset of expression of endogenous *npas4l*, resulted in upregulation of and ectopic expression of early hemato- vascular genes *etsrp* and *tal1* (Reischauer et al., 2016). We suspect that upregulation of *npas4l* target genes occurred because *arnt1* and *arnt2* are ubiquitously expressed at shield stage. Single cell RNA sequencing studies detected *arnt1* and *arnt2* mRNA throughout gastrulation, which encompasses shield stage, in zebrafish embryos (Baranasic et al., 2022). To test whether expression of *npas4l-*target genes requires *npas4l*, *arnt1,* and *arnt2*, we injected wild-type embryos at the 1-cell stage with either *npas4l* mRNA or *arnt1* and *arnt2* mRNA. The embryos were fixed at shield stage, and assayed for expression of *tal1* or *lmo2* using WISH. Wild-type embryos injected with *npas4l* mRNA exhibited robust expression of *tal1* compared to uninjected control embryos (**Figure 5A-B**). In contrast, embryos injected with *arnt1* and *arnt2* mRNA showed little or no expression of *tal1*, similar to uninjected embryos (**Figure 5C**). This suggests that neither *arnt1* nor *arnt2* can drive expression of *tal1* in the absence of *npas4l*. To directly test whether *arnt1* and *arnt2* can drive expression of *tal1* in the absence of *npas4l*, we injected *npas4l* mutant embryos with *npas4l* RNA or with *arnt1* and *arnt2* RNA (**Figure 5D-F)**.

**Figure 5.**
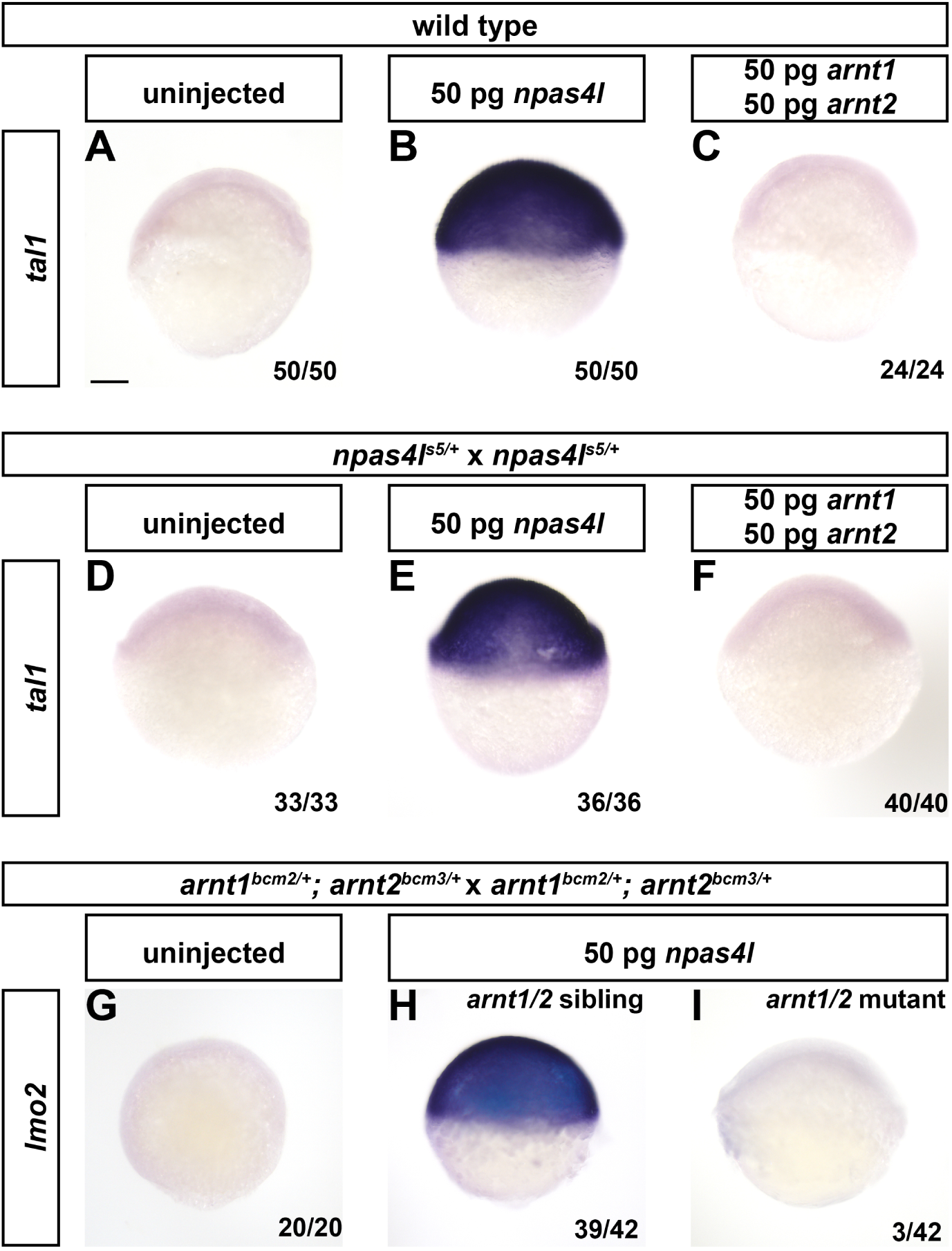
Expression of *npas4l*-target genes requires *npas4l*, *arnt1* and *arnt2*. Zebrafish embryos, derived from the indicted parental genotypes, were injected at the one-cell stage with 50 pg of *npas4l* mRNA or with 50 pg each of *arnt1* and *arnt2* mRNA (100 pg total RNA). Embryos were fixed at shield stage and probed for *tal1* or *lmo2, npas4l-*target genes, using whole mount in situ hybridization. Embryos were then genotyped. (A-C) In wild-type embryos, *tal1* is expressed following injection of *npas4l* RNA. *tal1* mRNA was not detected in uninjected embryos or following injection of *arnt1* and *arnt2* RNA. (D-F) Heterozygous *npas4l* mutant fish were crossed to each other and the resulting embryos were injected with the indicated RNA. *npas4l* RNA, but not *arnt1* and *arnt2* RNA, upregulated expression of *tal1*. Genotype of embryos in (E) n *=* 11 *npas4l^+/+^*, n = 15 *npas4l^s5/+^*, n = 10 *npas4l^s5/s5^*, (F) n = 12 *npas4l^+/+^*, n = 20 *npas4l^s5/+^*, n = 8 *npas4l^s5/s5^*. (G-I) *arnt1^bcm2/+^;arnt2^bcm3/+^* adults were crossed to each other and the resulting embryos were injected with *npas4l* RNA. Double homozygous embryos failed to upregulate *lmo2* (panel I) whereas sibling embryos upregulated *lmo2* (H). Genotype of embryos in (H) n = 1 *arnt1^+/+^;arnt2^+/+^*, n = 7 *arnt1^bcm2/+^;arnt2^bcm3/+^*, n = 1 *arnt1^+/+^;arnt2^bcm3/+^*, n = 8 *arnt1^bcm2/+^;arnt2^+/+^*, n= 4 *arnt1^+/+^;arnt2^bcm3/bcm3^*, n= 7 *arnt1^bcm2/bcm2^;arnt2^+/+^*, n = 7 *arnt1^bcm2/+^;arnt2^bcm3/bcm3^*, n = 4 *arnt1^bcm2/bcm2^;arnt2^bcm3/+^*, (I) n = 3 *arnt1^bcm2/bcm2^;arnt2^bcm3/bcm3^*. Embryos are oriented in a lateral view with the animal pole towards the top. Fractions in the bottom right corner indicate the number of embryos with the represented phenotype over the total number of embryos assayed. Scale bar in (A) applies to all panels, 100 μm.

Embryos injected with *npas4l* mRNA exhibited robust expression of *tal1* (**Figure 5E**), consistent with previously published results (Reischauer et al., 2016). However, all the embryos (derived from *npas4l^s5/+^* parents crossed to each other) injected with *arnt1* and *arnt2* mRNA showed little to no expression of *tal1* (**Figure 5F**). Since 25% of these embryos are *npas4l^s5/s5^*, we conclude that *arnt1* and *arnt2* cannot drive *tal1* expression in the absence of *npas4l*. Finally, to test if *npas4l* upregulates target genes in the absence of *arnt1* and *arnt2*, we injected *arnt1/2* mutant embryos with *npas4l* mRNA. We found that *arnt1/2* siblings showed upregulation of *npas4l-*target gene *lmo2*, but *arnt1/2* mutants did not show upregulation of *lmo2* (**Figure 5G-H**). From these results, we conclude that expression of *npas4l* and either *arnt1* or *arnt2* are required for upregulation of *npas4l*-target genes.

### Npas4l interacts with both Arnt1 and Arnt2

Npas4l is a Class I bHLH-PAS domain transcription factor (Reischauer et al., 2016), and as such, must heterodimerize with a Class II bHLH-PAS transcription factor, like Arnt1 and Arnt2, to regulate gene transcription. Considering that (1) bHLH-PAS transcription factors function as dimers, (2) *arnt1* and *arnt2* fail to upregulate hemato-vascular genes in the absence of *npas4l*, and (3) *npas4l* fails to upregulate hemato-vascular genes in *arnt1/2* mutants, therefore we hypothesize that Arnt1 and Arnt2 proteins form a transcription complex with Npas4l protein to regulate gene expression. To determine if Npas4l interacts with Arnt1 or Arnt2, we performed a coimmunoprecipitation experiment. Due to the absence of commercially available antibodies that recognize zebrafish Npas4l, Arnt1 or Arnt2 proteins, we co-expressed tagged versions of zebrafish *npas4l* (Npas4l-V5), *arnt1* (Arnt1-myc), and *arnt2* (Arnt2- HA) in HEK293T cells. We performed co-immunoprecipitation studies and found that Arnt1-myc interacts with Npas4l-V5 and that Arnt2-HA interacts with Npas4l-V5 (**Figure 6**, N= 2 biological replicates per condition). These results suggest that Npas4l uses either Arnt1 or Arnt2 or both as binding partners. Furthermore, these data indicate that Arnt1 and Arnt2 may be functionally interchangeable and redundant, at least in the case of Npas4l.

**Figure 6.**
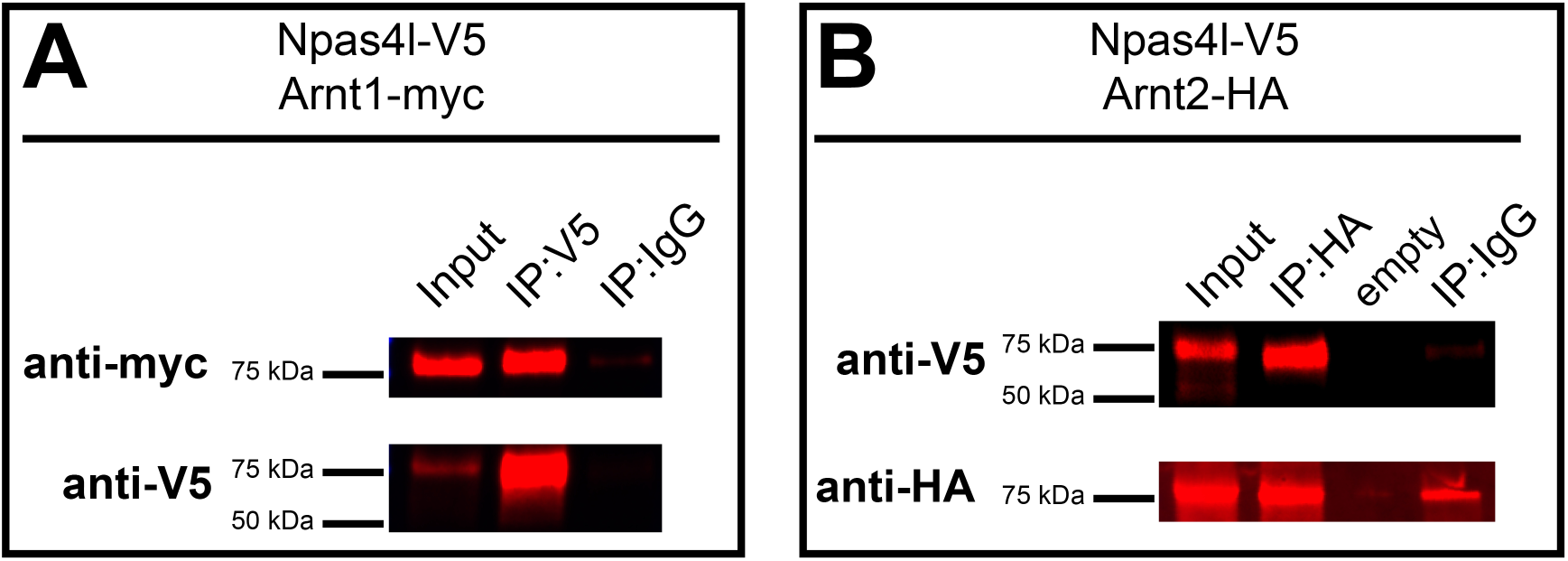
Arnt1 and Arnt2 co-immunoprecipitate with Npas4l. (A) HEK293T cells were transfected with zebrafish Npas4l-V5 and Arnt1-myc. Cells were homogenized (input) and immunoprecipitated (IP) with antibodies against V5 or an immunoglobulin G control (IgG). Samples were run on an immunoblot and probed using myc antibodies (anti-myc) to detect the presence of Arnt1-myc (top) or samples were probed using V5 antibodies (anti-V5) to verify successful pulldown of Npas4l-V5 (bottom). **(B)** HEK293T cells were transfected with zebrafish Npas4l-V5 and Arnt2-HA. Cells were homogenized and immunoprecipitated with antibodies against HA or an IgG control. Samples were run on an immunoblot and probed using V5 antibodies (anti-V5) to detect the presence of Npas4l-V5 (top) or samples were probed using HA antibodies (anti-HA) to verify successful pulldown of Arnt2-HA (bottom).

### Arnt1 and Arnt2 cannot functionally compensate for one another in the case of Ahr2

Our results demonstrate that Arnt1 and Arnt2 are functionally interchangeable and redundant when interacting with Npas4l in the context of hemato-vascular development. It is unclear if Arnt1 and Arnt2 are functionally interchangeable with all Class I bHLH-PAS domain transcription factors or just with Npas4l. To determine if *arnt1/arnt2* redundancy is unique to Npas4l, we examined if *arnt1* and *arnt2* are interchangeable with a different class I bHLH-PAS transcription factor, the aryl hydrocarbon receptor 2 (Ahr2). Previous studies demonstrate that Ahr2 can bind DNA in the presence of either Arnt1 or Arnt2 *in vitro* (Andreasen et al., 2002; Hirose et al., 1996; Tanguay et al., 2000), but whether Arnt1 and Arnt2 bind Ahr2 interchangeably *in vivo* is not known. Ahr2 is a ligand-dependent transcription factor. A known ligand of Ahr2 is 2,3,7,8-tetrachlorodibenzo-p-dioxin (TCDD), which we have previously shown induces cardiotoxicity in zebrafish larvae and requires *ahr2* (Souder and Gorelick, 2019). We asked whether TCDD causes cardiotoxicity in *arnt1* or *arnt2* single mutants, reasoning that if Ahr2 interacts with Arnt1 or Arnt2 interchangeably, then only *arnt1/2* double mutants, but not single mutants, would be resistant to TCDD exposure. We treated clutches of larvae from wild-type parents (N= 3 biological replicates, 10-60 larvae per clutch per drug treatment, 157 larvae total), *arnt1* heterozygous parents (N= 3-4 biological replicates per allele, 21-36 larvae per clutch per drug treatment, 533 larvae total, 248 larvae from allele *bcm1* and 285 larvae allele from *bcm2*), or *arnt2* heterozygous parents (N=3 biological replicates, 40-45 embryos per clutch per drug treatment, 253 larvae total) with either 0.1% DMSO (vehicle) or with 10 ng/mL TCDD from 1 dpf to 3 dpf **(Figure S4)**. Wild-type larvae exposed to TCDD exhibited pericardial edema and abnormal heart looping, consistent with previous results (**Figure 7A, D)**. *arnt2* mutants treated with TCDD exhibited a phenotype similar to wild-type (**Figure 7C, F)**. In contrast, *arnt1* mutants treated with TCDD showed normal heart looping and no pericardial edema, similar to vehicle-treated larvae (**Figure 7B, E)**. This demonstrates that *arnt1,* but not *arnt2,* is required for Ahr2-dependent TCDD toxicity. We conclude that *arnt1* and *arnt2* are not always functionally redundant *in vivo*. Certain Class I bHLH-PAS transcription factors may prefer to function with a specific Class II bHLH-PAS transcription faction.

**Figure 7.**
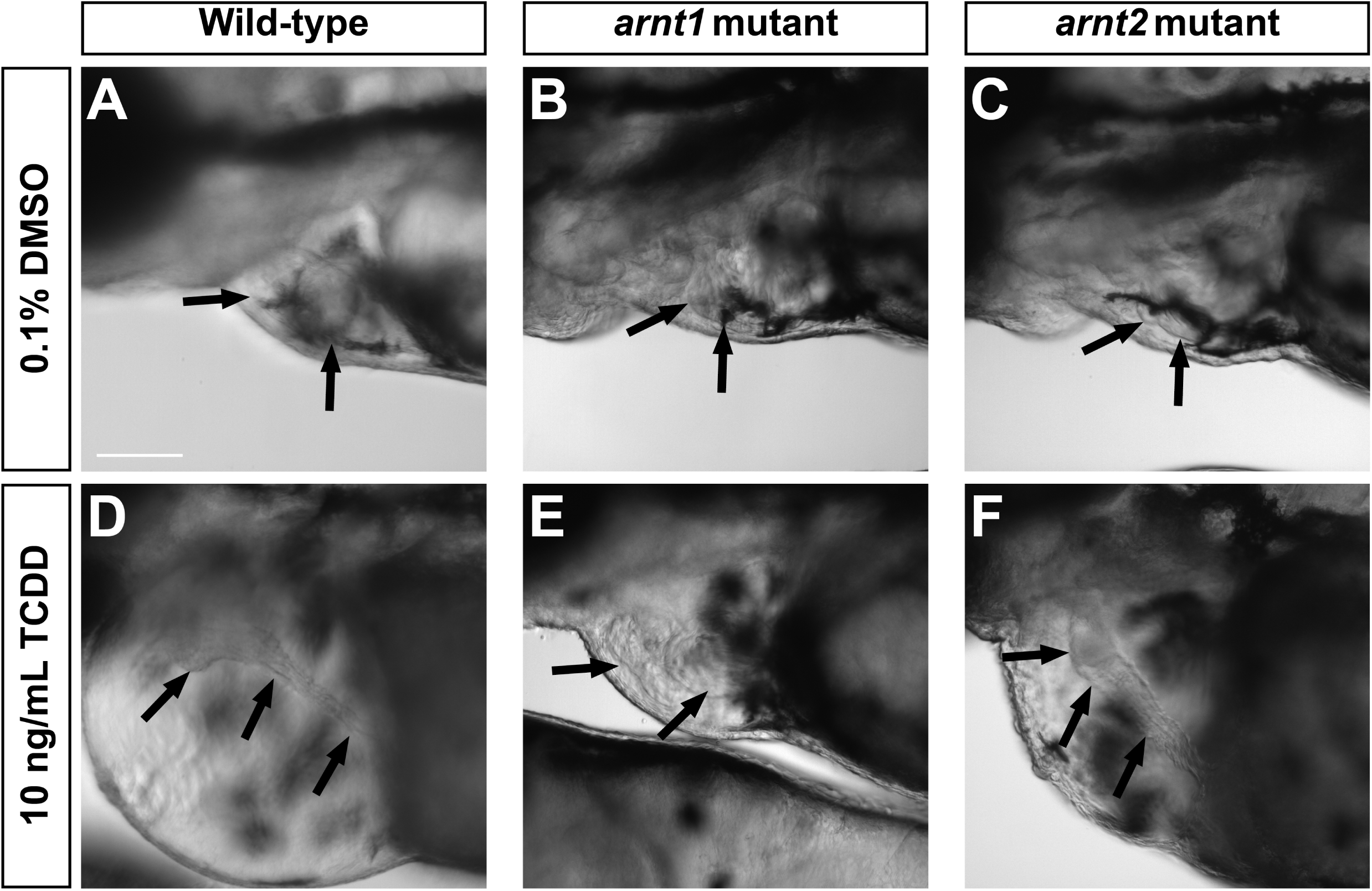
Mutation in *arnt1,* but not *arnt2,* prevents AHR2-dependent TCDD toxicity. Wild-type or mutant larvae were exposed to (**A, B, C**) 0.1% DMSO, or (**D, E, F**) 2,3,7,8-tetrachlorodibenzodioxin (TCDD) from 1-3 dpf and imaged at 3 dpf. (**A, D**) Wildtype larvae exposed to TCDD lack an S-shaped cardiac loop and exhibit cardiac edema, unlike vehicle treated wild-type siblings. (**B,E**) *arnt1* mutant larvae (*arnt1^bcm2/bcm2^*) treated with vehicle and TCDD show normal heart looping and do not have cardiac edema. (**C,F**) *arnt2* mutant larvae (*arnt2^bcm3/bcm3^*) treated with vehicle show expected heart looping similar to wild type siblings, but *arnt2* homozygous mutant larvae treated with TCDD lack an S- shaped cardiac loop similar to wild type and exhibit cardiac edema similar to TCDD treated wild-type larvae. Black arrows indicate location of the heart. Larvae were genotyped following imaging and representative images are shown. TCDD treated wild-type siblings of *arnt1* mutant larvae, n= 9 larvae across 3 clutches; TCDD treated *arnt1* mutant larvae, n=21 larvae across 3 clutches. TCDD treated wild-type siblings of *arnt2* larvae, n=19 larvae across 3 clutches; TCDD treated *arnt2* mutant larvae, n=22 larvae across 4 clutches. All larvae are oriented with anterior to the left and dorsal towards the top. Scale bar, 100 μm.

## DISCUSSION

We find that Arnt1 and Arnt2 are obligate coregulators of Npas4l. Together, *npas4l*, *arnt1* and *arnt2* genes regulate specification of hemato-vascular cells. *npas4l*, *arnt1* and *arnt2* are each necessary, but on their own not sufficient, for hemato-vascular specification. Npas4l serves as a master regulator, with Arnt proteins acting as permissive coregulators. Arnt1 and Arnt2 are functionally redundant in the case of Npas4l but are not functionally redundant in the case of Ahr2, where there is a preference for Ahr2- Arnt1 complex.

Class I bHLH-PAS proteins must dimerize with Class II bHLH-PAS proteins to transcribe their target genes (Card et al., 2005; Ema et al., 1996; Hirose et al., 1996; Jiang et al., 1996; Lees and Whitelaw, 1999; Michaud et al., 2000; Wu and Rastinejad, 2017). It has previously been established that Npas4l, a Class I bHLH-PAS transcription factor, is required for hemato-vascular development in zebrafish (Reischauer et al., 2016). To determine the Class II bHLH-PAS binding partner of Npas4l, we generated *arnt1* mutant and *arnt2* mutant zebrafish and expected one of these mutants to mimic the *npas4l* mutant phenotype. However, both *arnt1* mutant and *arnt2* single mutant larvae were viable to 5 dpf and were grossly indistinguishable from wild-type embryos. Only *arnt1/2* double mutant embryos exhibited a cardiovascular phenotype. Our results demonstrate that double homozygous *arnt1/2* mutant embryos have a similar phenotype to *npas4l* mutants (Reischauer et al., 2016; Stainier et al., 1995; Thompson et al., 1998) and that Npas4l can function with either Arnt1 or Arnt2. Thus, Arnt1 and Arnt2 act redundantly to specify hemato-vascular progenitor cells in zebrafish.

Are *arnt1* and *arnt2* functionally redundant with all Class I bHLH-PAS transcription factors? We argue no. In the case of the class I transcription factor Ahr2, both Arnt1 and Arnt2 bind Ahr2 *in vitro* (Lanham et al., 2011; Tanguay et al., 2000). However, we find that only Arnt1, and not Arnt2, is required for Ahr2-dependent cardiotoxicity in zebrafish embryos. This argues that *in vivo*, Ahr2 preferentially interacts with Arnt1 over Arnt2. Another class I protein, Sim1a, appears to preferentially interact with Arnt2 *in vivo*. *arnt2* mutant zebrafish embryos have fewer dopaminergic neurons in the brain compared to wild-type embryos (Löhr et al., 2009). The *arnt2* mutant phenotype is similar to the phenotype when Sim1a expression is knocked down (Löhr et al., 2009). These results suggest that in the case of the growth and/or survival of dopaminergic neurons, Sim1a preferentially interacts with Arnt2 versus Arnt1. To what degree this is due to differences in expression levels, versus an intrinsic structural preference for a particular protein-protein interaction, requires further study. Such interactions may also depend on the cell-type and developmental stage where the bHLH-PAS proteins function. Arnt1 and Arnt2 appear to be expressed ubiquitously during embryonic development, which suggests that such interactions may be due to factors other than expression levels.

Endothelial cell development and hematopoietic cell development are closely related during embryogenesis. Both cell types primarily arise from the lateral plate mesoderm (LPM) and share a common progenitor (Qian et al., 2005; Reischauer et al., 2016; Stainier et al., 1995; Thompson et al., 1998). Additionally, during definitive hematopoiesis, some HSCs are derived from specialized endothelial cells, called hemogenic endothelium (Boisset et al., 2010; Kissa and Herbomel, 2010). Both *npas4l* and *arnt1/2* mutants lack all blood cells but do have a small population of endothelial cells, which is restricted to the most posterior region of the animal (Figure 2 and (Reischauer et al., 2016)). In zebrafish, there is a population of endothelial cells in the caudal region of the embryo that are not derived from the LPM and differentiate in the absence of *npas4l* expression (Pak et al., 2020). We hypothesize that *arnt1* and *arnt2* are required for the differentiation of endothelial cells from the LPM but are not required for the differentiation of endothelial cells, such as the caudal population, from outside the LPM. The transcriptomic signature of the caudal population of endothelial cells was distinct from endothelial cells that require *npas4l* for differentiation, with genes for somitogenesis and neurogenesis highly enriched compared to LPM-derived endothelial cells (Pak et al., 2020). Based on this transcriptome analysis, we speculate that the caudal population of endothelial cells fails to express genes required for hemogenic endothelium and that no blood cells are derived from the caudal population of endothelial cells. This is consistent with the observation that the caudal population of endothelial cells, but not blood cells, are present in *npas4l* or *arnt1/2* mutants.

Arnt proteins interact with multiple class I bHLH-PAS domain proteins to regulate cardiovascular development. Our results focused on the role of Arnt proteins in hemato-vascular specification, however Arnt proteins are likely involved in later steps of endothelial cell development. Differentiation of hemogenic endothelium in zebrafish and in mice is dependent upon several Class I bHLH-PAS transcription factors: Hypoxia inducible factor 1 alpha (mouse *Hif1α*, zebrafish genes *hif1aa*, *hif1ab)* and hypoxia inducible factor 2 alpha (mouse *Hif2α,* zebrafish genes *epas1a*, *epas1b*) (Gerri et al., 2018; Ramírez-Bergeron et al., 2004). Given that these HIF proteins are Class I bHLH-PAS transcription factors, it is likely that one or both Arnt proteins are required for HIF-dependent differentiation of hemogenic endothelium.

The function of Arnt proteins in zebrafish cardiovascular development is likely conserved in mice. Arnt1 is essential for the viability of adult hematopoietic stem cells in mice and is required for the survival of hematopoietic progenitor cells in the fetal mouse liver (Krock et al., 2015). Homozygous *Arnt1* mutant mice are not viable beyond E10.5 and show severe defects in vascularization, suggesting that Arnt1 is essential for normal vascular development (Kozak et al., 1997; Maltepe et al., 1997). The degree to which Arnt2 contributes to hemato-vascular development in mice is not well-understood. Arnt2 has primarily been studied within the context of neural development (Hao et al., 2013; Hosoya et al., 2001; Jain et al., 1998), however, there is some evidence to suggest that *Arnt1* and *Arnt2* may exhibit functional redundancy during development of non-neural tissues (Keith et al., 2001; Sekine et al., 2006). We observed lower than expected numbers of *arnt1^bcm2/bcm2^; arnt2^bcm3/+^* adult zebrafish derived from *arnt1^bcm2/+^; arnt2^bcm3/+^* parents. This suggests that *arnt1* and *arnt2* have a dose-dependent effect on the survival or fitness of the animal, where anything less than four wild-type alleles (two *arnt1* alleles, two *arnt2* alleles) is deleterious. A similar dose-dependent effect on survival can be seen in mouse embryos from *Arnt1^+/-^;Arnt2^+/-^* parents (Keith et al., 2001). The mechanisms behind these dose- dependent gene effects are not known.

While zebrafish *arnt1* and *arnt2* genes are conserved in mammals, there is no known equivalent of zebrafish *npas4l* in mammalian genomes. Zebrafish Npas4l shows the highest sequence homology to mouse Npas4 (Reischauer et al., 2016). Although Npas4 is known to dimerize with Arnt1 and Arnt2 *in vivo* (Brigidi et al., 2019), mammalian Npas4 is unlikely to be the functional equivalent to zebrafish Npas4l because homozygous Npas4 mutant mice are viable to adulthood and do not show any overt vascular or hematopoietic defects (Bloodgood et al., 2013). Given our results that *npas4l* requires *arnt1* or *arnt2* to act as co-regulators of hemato-vascular specification, and the abundance of conserved genes that regulate vascular development in zebrafish and humans (Hogan and Schulte-Merker, 2017), we speculate that the functional mammalian equivalent of *npas4l* is a Class I bHLH-PAS transcription factor, or multiple such factors, that function together with Arnt1 or Arnt2.

## Acknowledgements

We thank Lauren Pandolfo and the staff of the Baylor College of Medicine aquatics facility for taking care of our zebrafish colony, Rosa Uribe and Adam Howard from Rice University for help with WIHCR, and Paula Pimienta-Ramirez and Yunping Lei in the CPEH DNA sequencing core at BCM for help with sequencing embryos following in situ hybridization. This work was supported by NIH grants R01ES026337 (to DAG), P30ES030285 (pilot award to DAG), and T32DK060445 (to MJE).

## Supplemental files

Supplemental Table 1. Oligonucleotide primers and probes used in this study

Supplemental Table 2. Genotype versus phenotype correlation of embryos probed for *etsrp* (related to Figure 3)

Supplemental Table 3. Genotype versus phenotype correlation of embryos probed for *tal1* (related to Figure 3)

Supplemental Table 4. Genotype versus phenotype correlation of embryos probed for *lmo2* (related to Figure 3)

Supplemental Table 5. Genotype versus phenotype correlation of embryos probed for *egfl7* (related to Supplemental Figure 1)

Supplemental Table 6. Genotype versus phenotype correlation of embryos probed for *npas4l* (related to Supplemental Figure 2)

Supplemental Figure 1. a*r*nt1*/2* mutants show reduced expression of *egfl7* in the lateral plate mesoderm (related to Figure 3)

Supplemental Figure 2. a*r*nt1*/2* mutants and *arnt1/2* siblings have normal expression of *npas4l* (related to Figure 3)

Supplemental Figure 3. n*p*as4l and *arnt2* mRNA, but not *arnt1*, are enriched in the posterior lateral plate mesoderm (related to Figure 4)

Supplemental Figure 4. a*r*nt1 mutants, but not *arnt2* mutants, are resistant to TCDD toxicity (related to Figure 7)

Movie 1.mp4 *arnt1^bcm2/+^;arnt2^bcm3/+^* 50 hpf, head and heart

Movie 2.mp4 *arnt1^bcm2/+^;arnt2^bcm3/+^* 50 hpf, blood flow in tail

Movie 3.mp4 *arnt1^bcm2/bcm2^;arnt2^bcm3/bcm3^* 50 hpf, head and heart

Movie 4.mp4 *arnt1^bcm2/bcm2^;arnt2^bcm3/bcm3^* 50 hpf, blood flow in tail

Movie 5.mp4 *arnt1^bcm2/+^;arnt2^bcm3/+^* 2 dpf, head and heart

Movie 6.mp4 *arnt1^bcm2/+^;arnt2^bcm3/+^* 2 dpf, blood flow in tail

Movie 7.mp4 *arnt1^bcm2/bcm2^;arnt2^bcm3/bcm3^* 2 dpf, head and heart

Movie 8.mp4 *arnt1^bcm2/bcm2^;arnt2^bcm3/bcm3^* 2 dpf, blood flow in tail

Movies 1 and 2 from the same embryo, movies 3 and 4 from the same embryo, movies 5 and 6 from the same embryo, movies 7 and 8 from the same embryo.

---

**Table S1.**
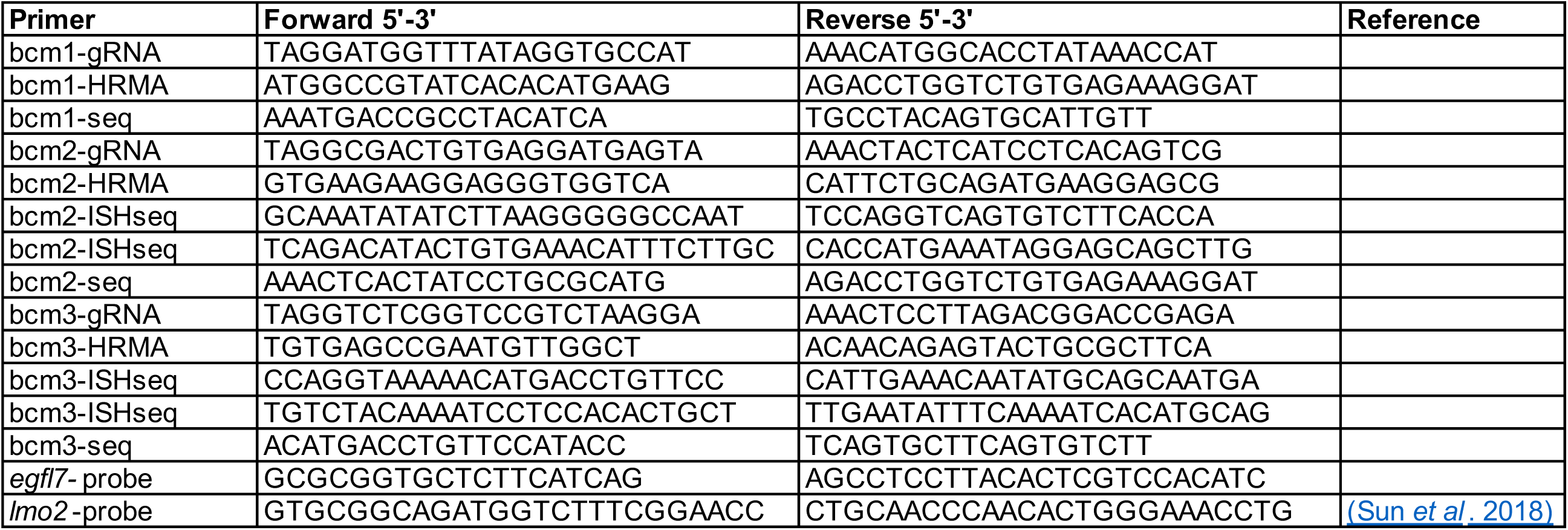
Oligonucleotide primers and probes used in this study

**Table S2.** Genotype versus phenotype correlation of embryos probed for *etsrp* (See Figure 3). There was a significant correlation between the number of mutant alleles and *etsrp* expression phenotype, *r(47)= - 0.9, p<0.001*

| Genotype |  | Phenotype (number of embryos) |  |
| --- | --- | --- | --- |
| Allele designation | Number of mutant alleles | Normal expression | Reduced expression |
| *arnt1*<sup>bcm2/bcm2</sup> ; *arnt2*<sup>bcm3/bcm3</sup> | 4 | 2 | 24 |
| *arnt1*<sup>bcm2/bcm2</sup> ; *arnt2*<sup>bcm3/+</sup> | 3 | 4 | 0 |
| *arnt1*<sup>bcm2/+</sup> ; *arnt2*<sup>bcm3/bcm3</sup> |  | 4 | 0 |
| *arnt1*<sup>bcm2/+</sup> ; *arnt2*<sup>bcm3/+</sup> | 2 | 5 | 0 |
| *arnt1*<sup>bcm2/bcm2</sup> |  | 3 | 0 |
| *arnt2*<sup>bcm3/bcm3</sup> |  | 2 | 0 |
| *arnt1*<sup>bcm2/+</sup> | 1 | 4 | 0 |
| *arnt2*<sup>bcm3/+</sup> |  | 3 | 0 |
| wild-type | 0 | 1 | 0 |

**Table S3.** Genotype versus phenotype correlation of embryos probed for tal1 (See Figure 3). There was a significant correlation between the number of mutant alleles and tal1 expression phenotype, r(25)= -0.6, p<0.001

| Genotype |  | Phenotype (number of embryos) |  |
| --- | --- | --- | --- |
| Allele designation | Number of mutant alleles | Normal expression | Reduced expression |
| *arnt1*<sup>bcm2/bcm2</sup> ; *arnt2*<sup>bcm3/bcm3</sup> | 4 | 1 | 14 |
| *arnt1*<sup>bcm2/bcm2</sup> ; *arnt2*<sup>bcm3/+</sup> | 3 | 3 | 3 |
| *arnt1*<sup>bcm2/+</sup> ; *arnt2*<sup>bcm3/bcm3</sup> |  | 1 | 0 |
| *arnt1*<sup>bcm2/+</sup> ; *arnt2*<sup>bcm3/+</sup> | 2 | 1 | 0 |
| *arnt1*<sup>bcm2/bcm2</sup> |  | 0 | 0 |
| *arnt2*<sup>bcm3/bcm3</sup> |  | 0 | 0 |
| *arnt1*<sup>bcm2/+</sup> | 1 | 1 | 0 |
| *arnt2*<sup>bcm3/+</sup> |  | 2 | 0 |
| wild-type | 0 | 0 | 0 |

**Table S4.** Genotype versus phenotype correlation of embryos probed for *lmo2* (See Figure 3). There was a significant correlation between the number of mutant alleles and *lmo2* expression phenotype, *r(184)= -0.2, p<0.005*

| Genotype |  | Phenotype (number of embryos) |  |
| --- | --- | --- | --- |
| Allele designation | Number of mutant alleles | Normal expression | Reduced expression |
| *arnt1*<sup>bcm2/bcm2</sup> ; *arnt2*<sup>bcm3/bcm3</sup> | 4 | 5 | 7 |
| *arnt1*<sup>bcm2/bcm2</sup> ; *arnt2*<sup>bcm3/+</sup> | 3 | 23 | 5 |
| *arnt1*<sup>bcm2/+</sup> ; *arnt2*<sup>bcm3/bcm3</sup> |  | 13 | 3 |
| *arnt1*<sup>bcm2/+</sup> ; *arnt2*<sup>bcm3/+</sup> | 2 | 47 | 3 |
| *arnt1*<sup>bcm2/bcm2</sup> |  | 5 | 0 |
| *arnt2*<sup>bcm3/bcm3</sup> |  | 10 | 2 |
| *arnt1*<sup>bcm2/+</sup> | 1 | 28 | 1 |
| *arnt2*<sup>bcm3/+</sup> |  | 19 | 2 |
| wild-type | 0 | 7 | 2 |

**Table S5.**
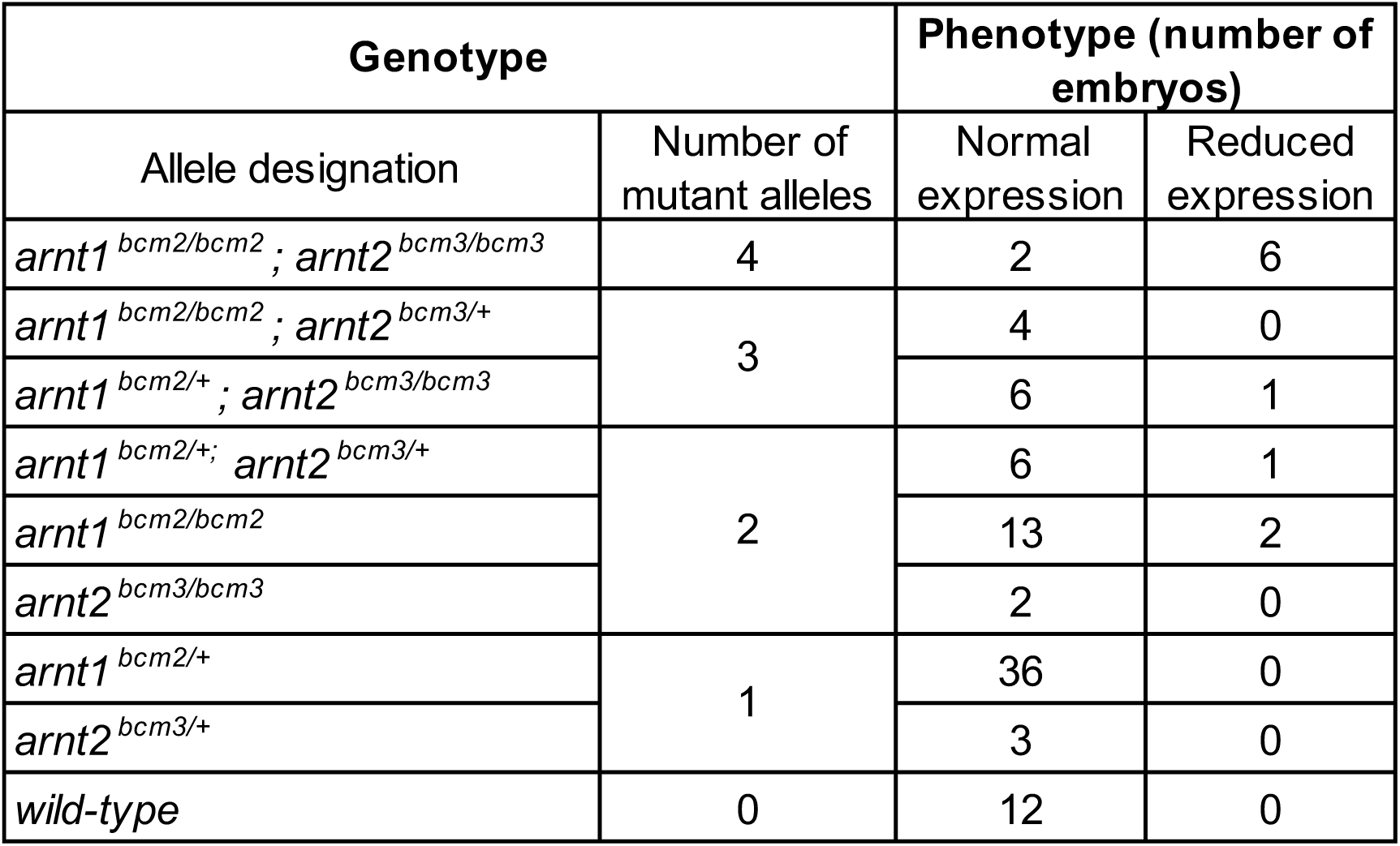
Genotype versus phenotype correlation of embryos probed for egfl7 (See Supplemental Figure 1). There was a significant correlation between the number of mutant alleles and egfl7 expression phenotype, r(94)= -0.5, p<0.001

**Table S6.** Genotype versusphenotype correlation of embryos probed for *npas4l* (See Supplemental Figure 2). There was no significant correlation between the number of mutant alleles and *npas4l* expression phenotype, *r(69)=-0.05, p>0.05*

| Genotype |  | Phenotype (number of embryos) |  |
| --- | --- | --- | --- |
| Allele designation | Number of mutant alleles | Normal expression | Reduced expression |
| *arnt1*<sup>bcm2/bcm2</sup> ; *arnt2*<sup>bcm3/bcm3</sup> | 4 | 2 | 0 |
| *arnt1*<sup>bcm2/bcm2</sup> ; *arnt2*<sup>bcm3/+</sup> | 3 | 4 | 0 |
| *arnt1*<sup>bcm2/+</sup> ; *arnt2*<sup>bcm3/bcm3</sup> |  | 11 | 0 |
| *arnt1*<sup>bcm2/+</sup> ; *arnt2*<sup>bcm3/+</sup> | 2 | 19 | 1 |
| *arnt1*<sup>bcm2/bcm2</sup> |  | 3 | 1 |
| *arnt2*<sup>bcm3/bcm3</sup> |  | 5 | 0 |
| *arnt1*<sup>bcm2/+</sup> | 1 | 11 | 0 |
| *arnt2*<sup>bcm3/+</sup> |  | 5 | 0 |
| wild-type | 0 | 8 | 0 |

**Supplemental Figure 1.**
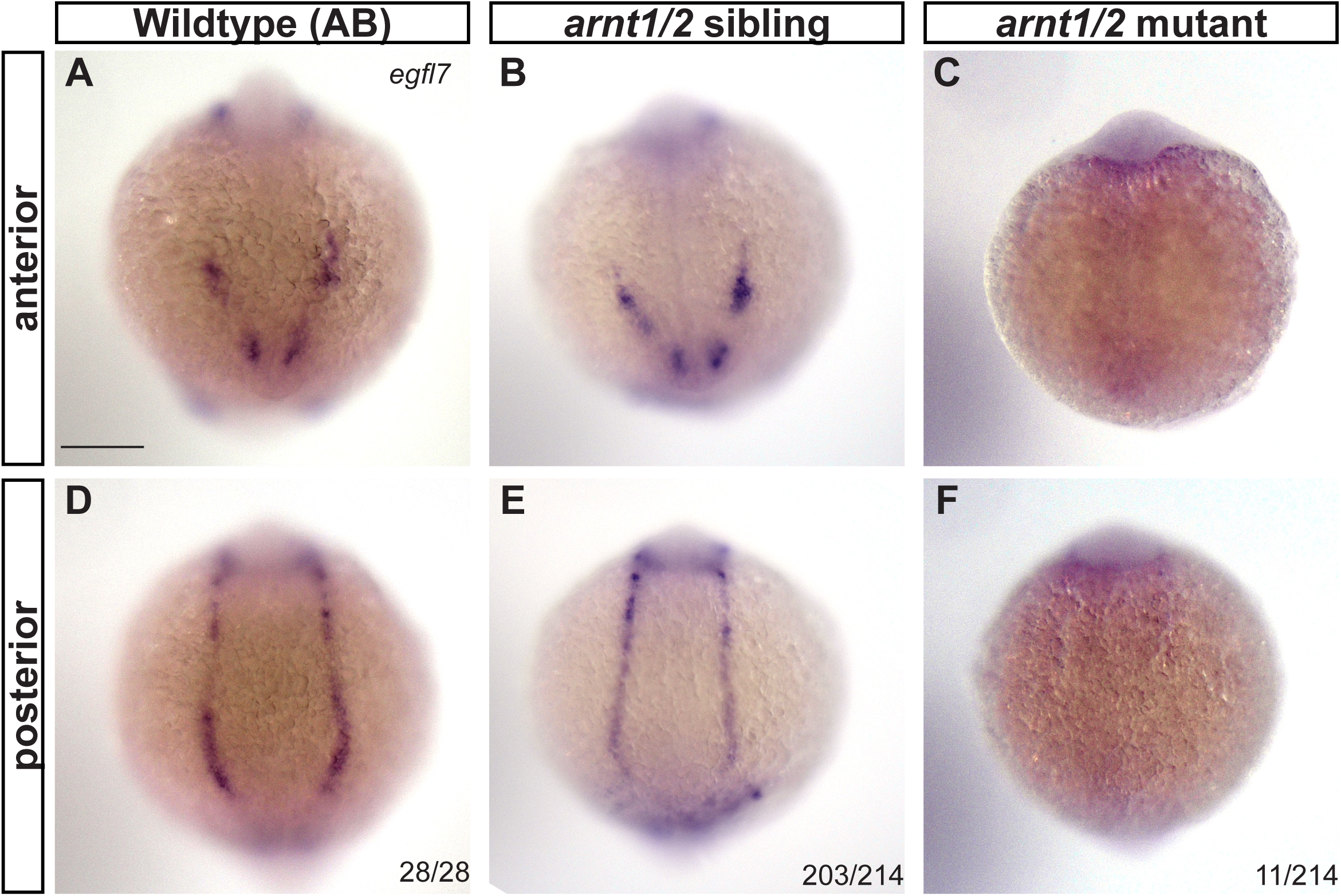
a*r*nt1*/2* mutants show reduced expression of *egfl7* in the lateral plate mesoderm. Whole-mount in situ hybridization for *egfl7* was performed on 10 somite stage embryos from *arnt1^bcm2/+^;arnt2^bcm3/+^* parents (N=2 biological replicates, 94-120 embryos per clutch, 214 total embryos) or from wildtype (AB) parents (N=1 biological replicate, 28 total embryos). (**A** and **D**) expression of *egfl7* in wildtype embryos is restricted to the anterior lateral plate mesoderm (ALPM) (**A**) and the posterior lateral plate mesoderm (PLPM) (**D**). (**B** and **E**) Expression of *egfl7* was observed in the ALPM (**B**) and the PLPM (**E**) in 95% of embryos from *arnt1^bcm2/+^; arnt2^bcm3/+^* parents (n=203 out of 214 total embryos). (**C** and **F**) Limited or no expression of *egfl7* was observed in the ALPM and PLPM in 5% of embryos from *arnt1^bcm2/+^;arnt2^bcm3/+^* parents (n=11 out of 214 total embryos), which is consistent with the expected Mendelian ratio of 6.25% *arnt1^bcm2/bcm2^;arnt2^bcm3/bcm3^* (*arnt1/2* mutants, binomial test, p<0.05). All embryos are oriented with dorsal towards the top of the image. Fractions in bottom right corners refer to the number of embryos with the indicated phenotype over the total number of embryos examined. See **Supplemental Table 5** for genotype frequencies. Scale bar, 100μm.

**Supplemental Figure 2.**
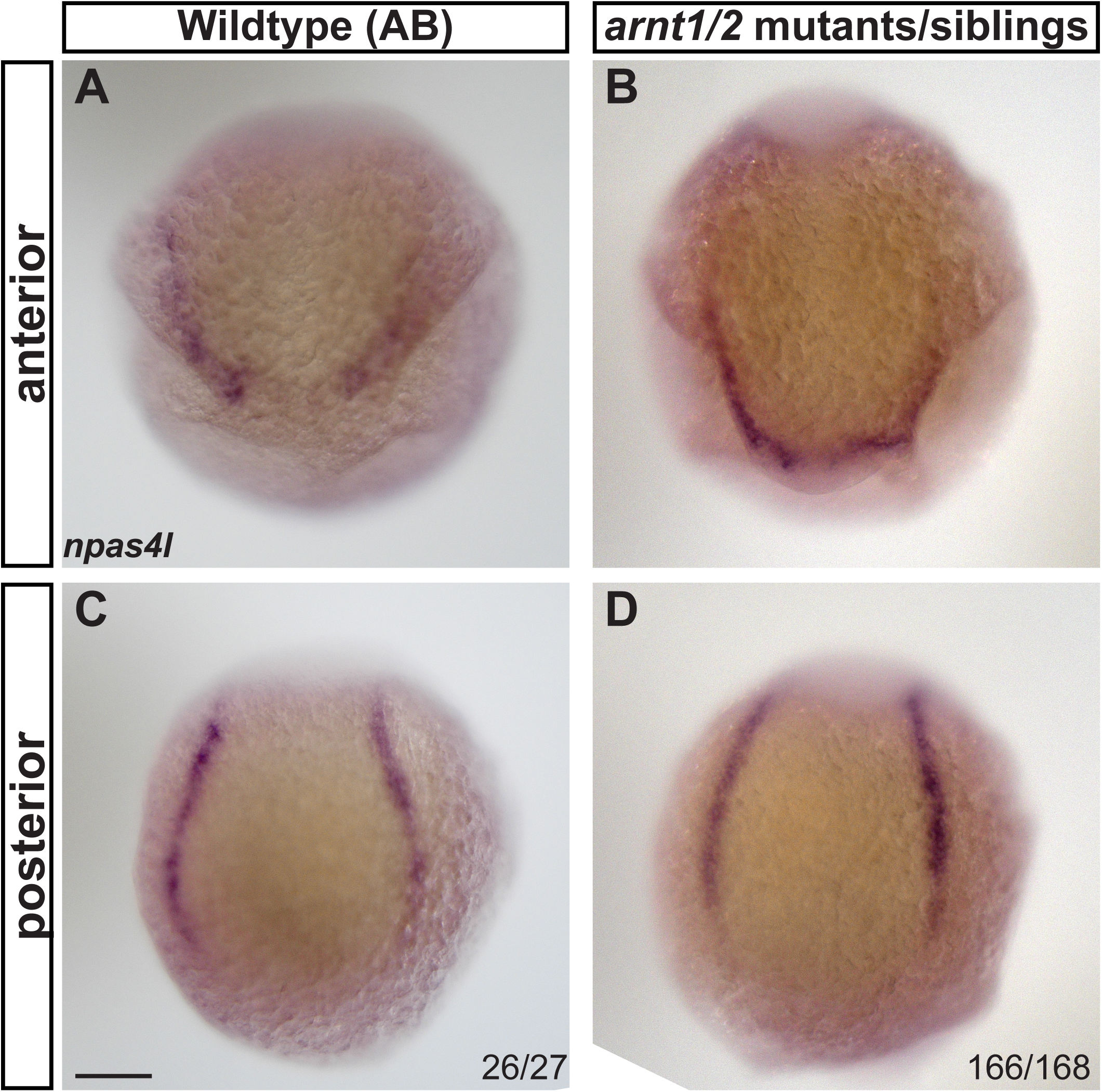
a*r*nt1*/2* mutants and *arnt1/2* siblings have normal expression of *npas4l.* Whole-mount in situ hybridization for *npas4l* on 4 somite stage embryos from *arnt1^bcm2/+^; arnt2^bcm3/+^* parents (N=2 biolgical replicates, 68-100 embryos per clutch, n= 168 embryos total) or from wild-type (AB) parents (N=1 biological replicate, n=27 embryos total). (**A**) Expression of *npas4l* was observed in the anterior lateral plate mesoderm (ALPM) of wild-type embryos. (**B**) Expression of *npas4l w*as observed in the ALPM embryos from *arnt1^bcm2/+^; arnt2^bcm3/+^* parents, including the *arnt1/2* mutants. (**C**) Expression of *npas4l* was observed in the PLPM of wild-type embryos. (**D**) Expression of *npas4l w*as observed in the posterior lateral plate meosderm (PLPM) embryos from *arnt1^bcm2/+^; arnt2^bcm3/+^* parents, including the *arnt1/2* mutants. Embryos oriented with dorsal towards the top of the image. The fractions in the right-hand corners of **C** and **D** refer to the number of embryos with the represented phenotypes over the total number of embryos. Scale bar, 100 μm.

**Supplemental Figure 3.**
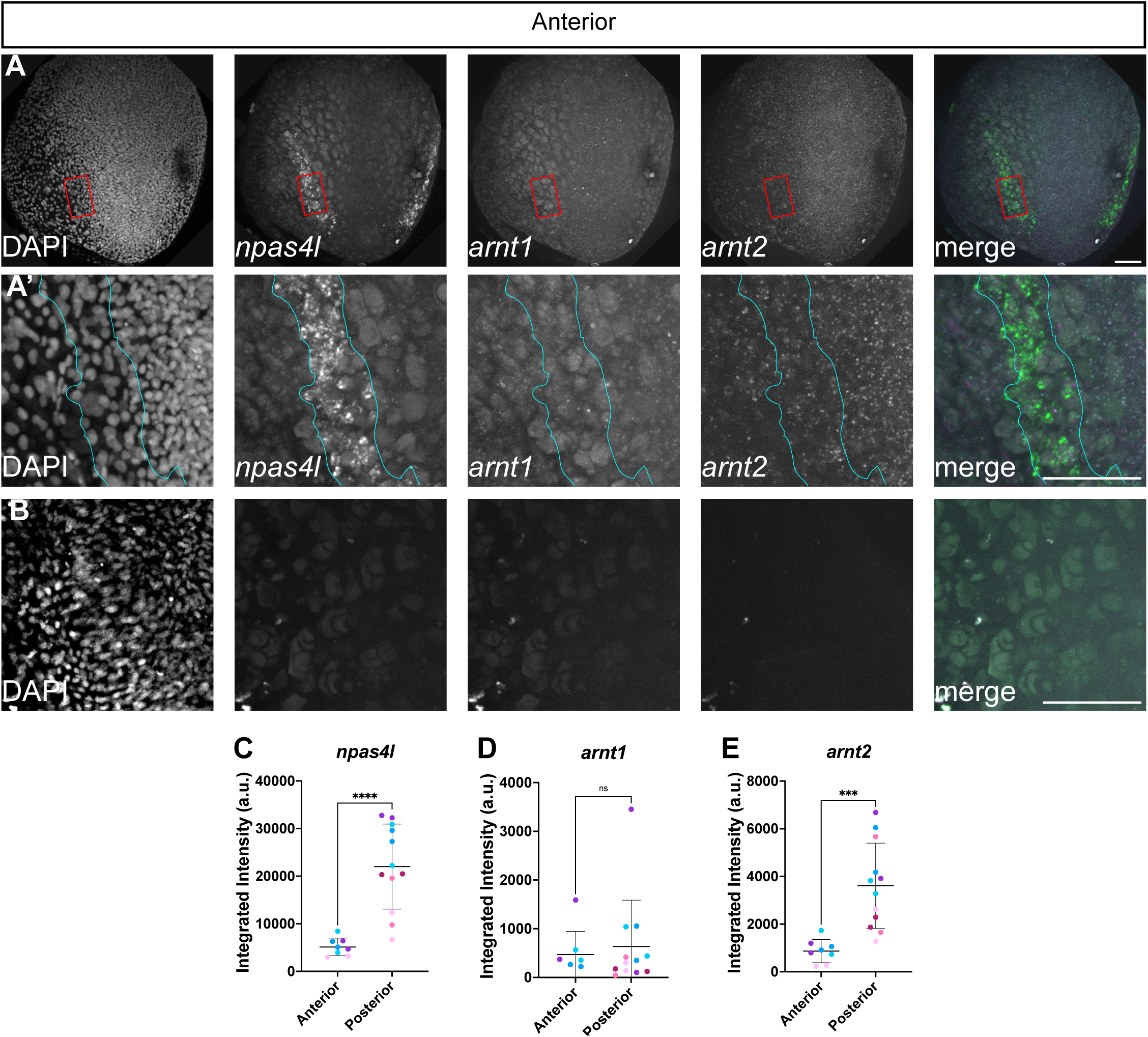
Expression of *npas4l*, *arnt1* and *arnt2* mRNA in the anterior lateral plate mesoderm. Companion to Figure 4, which shows expression in the posterior lateral plate mesoderm. Whole-mount *in situ* hybridization chain reaction was performed on wild-type (AB) embryos at the 4 somite stage. **(A)** Embryos were probed for *npas4l, arnt1,* and *arnt2* expression and counter-stained with DAPI. **(A’)** Higher magnification views of the boxed areas in (A) shows the anterior lateral plate mesoderm, outlined in cyan. N=3 biological replicates, 5-9 embryos per replicate, 22 embryos total. **(B)** To demonstrate that the observed labeling is not autofluorescence of the embryo or the yolk, wild-type embryos were processed identically to those in (A), but no anti-sense RNA probe sets were added. The views in (B) are identical magnification as those in (A’). N=3 biological replicates, 6-12 embryos per replicate, 28 embryos total. Images are maximum intensity projections from z-stacks taken every 2.5 μm. In merged panels, *npas4l* is green, *arnt1* is magenta, *arnt2* is cyan. The anterior portion of the embryo is in view, dorsal is towards the top of the image. Scale bars = 100 μm. **(C-E)** *npas4l* and *arnt2* mRNA, but not *arnt1*, are enriched in the posterior lateral plate mesoderm. Quantification of the fluorescence integrated intensity of *npas4l, arnt1,* and *arnt2* labeling within the *npas4l*-positive cells of the anterior and posterior lateral plate mesoderm (see panel A and Figure 4). We calculated the integrated intensity of fluorescence (pixel area, pixel intensity) over each side of the anterior or posterior LPM in maximum intensity projections. Each point represents the integrated intensity from one side of the LPM from a single embryo (right or left side), where each embryo has two bilateral *npas4l*-positive regions of the LPM. Points of the same color are measurements from opposite sides of the LPM from the same embryo. 6 representative embryos were used for this analysis, in 2 of these embryos the anterior LPM was damaged during imaging and could not be analyzed. **(C)** *npas4l* expression was statistically significantly increased in the posterior versus anterior LPM, Welch’s t-test, *t(12.36)=6.4, p<0.0001.* **(D)** *arnt1* was expressed at similar levels in the anterior and posterior, Mann-Whitney, U=45, p>0.05. **(E)** *arnt2* expression was statistically significantly increased in the posterior versus anterior LPM, Welch’s t-test, *t(11)= 5.0, p<0.0005*.

**Supplemental Figure 4.**
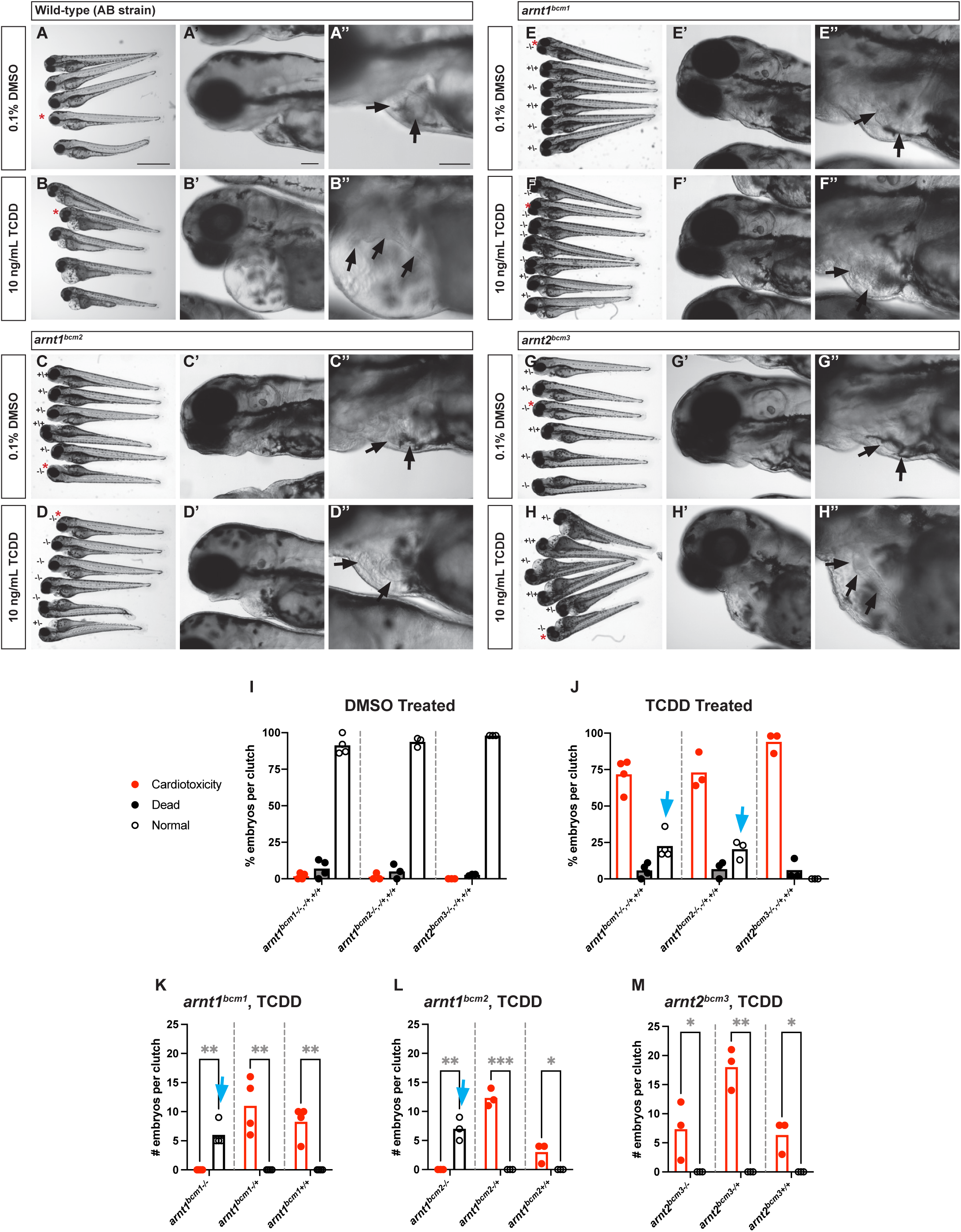
a*r*nt1 mutants, but not *arnt2* mutants, are resistant to TCDD toxicity. Representative images of larvae at 3 days post fertilization (dpf) exposed to either DMSO or TCDD. Mutant larvae were derived from heterozygous parents and genotyped following imaging (denoted as wild-type +/+, heterozygous -/+, or homozygous-/- in panels **C-H**). (**A-H**) overview images of treated larvae, red asterisks denote larvae detailed in panels A’-H’ and A”-H”. Black arrows in panels A”-H” point to the heart within the pericardial cavity. Pericardial edema and incomplete heart looping is present in 100% of wild-type (n=3 biological replicates per genotype, 10-60 larvae per clutch per treatment, 157 larvae total) and *arnt2* mutant larvae (n=3 biological replicates per genotype, 40-50 larvae per clutch per treatment, 253 larvae total). In contrast, 100% of *arnt1* mutants (3-4 clutches per allele, 21-36 larvae per clutch per treatment, 533 larvae total, 248 larvae from allele bcm1 and 285 larvae allele from bcm2) are resistant to TCDD cardiotoxicity. Scale bars, 100 μm. (**I** and **J**) Percent larvae from each clutch demonstrating cardiotoxicity (red dots/red bars), mortality (black dots/gray bars), or normal development (white dots/white bars) at 3 dpf. All clutches were derived from heterozygous parents and embryos were analyzed prior to genotyping. Each clutch contain a mix of wild-type, heterozygous, and homozygous larvae. Each dot represents the percent of larvae from a single clutch and bars represent the mean. (**J**) In clutches derived from *arnt2^bcm3/+^* parents, nearly 100% of embryos exhibited cardiotoxicity following TCDD exposure. In clutches derived from *arnt1* heterozygous parents, approximately 75% of embryos exhibited cardiotoxicity or death, while 25% of embryos exhibited a normal phenotype (blue arrows), consistent with the expected Mendelian ratio of *arnt1* homozygotes. Normal larvae were not observed in wild-type or *arnt2* mixed clutches exposed to TCDD. (**K-M**) We genotyped the embryos from (**J**) and graphed the number of embryos per clutch exhibiting cardiotoxicity (red dots/red bars) or normal development (white dots/white bars). Each dot represents the number of larvae from a single clutch. (**K,L**) All *arnt1* homozygous mutants (*bcm1* or *bcm2*) exhibited normal phenotype following TCDD exposure (blue arrows), while *arnt1* heterozygotes and (**M**) *arnt2* mutants exhibited cardiotoxicity. Unpaired Welch’s t-tests, * *p<*0.05, ** *p<*0.005, and *** *p<*0.001.

